# Shared gene targets of the ATF4 and p53 transcriptional networks

**DOI:** 10.1101/2023.03.15.532778

**Authors:** Gabriele Baniulyte, Serene A. Durham, Lauren E. Merchant, Morgan A. Sammons

## Abstract

The master tumor suppressor p53 regulates multiple cell fate decisions, like cell cycle arrest and apoptosis, via transcriptional control of a broad gene network. Dysfunction in the p53 network is common in cancer, often through mutations that inactivate p53 or other members of the pathway. Induction of tumor-specific cell death by restoration of p53 activity without off-target effects has gained significant interest in the field. In this study, we explore the gene regulatory mechanisms underlying a putative anti-cancer strategy involving stimulation of the p53-independent Integrated Stress Response (ISR). Our data demonstrate the p53 and ISR pathways converge to independently regulate common metabolic and pro-apoptotic genes. We investigated the architecture of multiple gene regulatory elements bound by p53 and the ISR effector ATF4 controlling this shared regulation. We identified additional key transcription factors that control basal and stress-induced regulation of these shared p53 and ATF4 target genes. Thus, our results provide significant new molecular and genetic insight into gene regulatory networks and transcription factors that are the target of numerous antitumor therapies.

## Introduction

The global rewiring of cellular anabolic and catabolic processes that result from homeostatic changes include dynamic control of RNA and protein synthesis and turnover (Vihervaara et al., 2018; Advani and Ivanov, 2019). The DNA damage-inducible transcription factor, p53, directly activates transcription of a broad range of target genes involved in DNA repair, cell cycle arrest, and apoptosis. The well-described tumor suppressor function of p53 primarily relies on transcriptional activation of these target genes and their ability to mitigate the consequences of damaged DNA. Nearly half of human malignancies harbor mutations in p53 that facilitate and promote metastasis, tumorigenesis, and resistance to apoptosis (Zhu et al., 2015; Mantovani et al., 2019). These mutations generally lead to loss of DNA binding and an inability to transactivate canonical anti-proliferative p53 target genes (Bykov et al., 2018). Genotoxic chemotherapeutics, like doxorubicin and etoposide, are clinically relevant activators of wild-type p53, but the potential risk of resistance and secondary malignancies due to increased mutational burden remains a significant concern (Aziz et al., 2011). Given the powerful tumor suppression abilities of p53, restoration of the p53-regulated transcriptome without inducing additional DNA damage represents an intriguing approach for development of anticancer strategies and therapeutics.

Non-genotoxic, small molecule activation of the p53 pathway has been proposed as a potential solution. The first general approach involves small-molecule targeting of mutant p53 to restore its wild-type function or prevent dominant-negative/gain-of-function activities (Yu et al., 2012; Chowdhury et al., 2014; Zhang et al., 2015; Bykov et al., 2018). A second approach uses compounds like the MDM2 inhibitor nutlin-3A to activate wild-type p53 in a non-genotoxic fashion, although early clinical trials suggest these approaches have limited efficacy when used alone (Andreeff et al., 2016; Kocik et al., 2019; Montesinos et al., 2020). A third approach involves bypassing p53 altogether via compounds that activate key anti-proliferative p53 targets in p53-deficient tumors (Hernandez Borrero et al., 2021; Tian et al., 2021). These compounds engage the Integrated Stress Response (ISR) network which is an effector of anti-proliferative and cell death gene expression programs. Interestingly, simultaneous ISR activation and MDM2 inhibition led to significant cell death and tumor regression not observed when the approaches were used individually (Andrysik et al., 2022), suggesting these pathways may work synergistically. These new ISR-stimulating approaches may be broadly applicable, as wild-type p53, p53-deficient, and p53 missense mutation-containing tumors could all be targeted. Thus, further exploration into the genetic and biochemical basis underlying this shared synergy between the p53 and ISR gene regulatory networks is needed for the design of more efficacious therapeutics with known mechanisms of action.

Here, we examine shared gene targets and regulatory strategies of two distinct stress-dependent pathways, the p53 gene regulatory network (GRN) and the ATF4-dependent Integrated Stress Response (ISR) pathway. We identify an enhancer element required for p53-dependent induction of *ATF3* in response to DNA damage, however, is not directly required for transcription of *ATF3* under ISR activating conditions. ISR-dependent induction of *ATF3* requires ATF4 via binding and regulation of the *ATF3* promoter. We identified a second regulatory strategy whereby ATF4 and p53 target a shared enhancer to control *GADD45A,* another common gene target. Our data suggest that the p53 and ISR pathways have shared gene targets and begin to unravel the DNA encoding and TF requirements for engagement of *cis*-regulatory elements driving these behaviors.

## Materials & Methods

### Cell Culture and Treatments

The human colorectal cancer cell lines, HCT116 TP53^+/+^ and HCT116 TP53^-/-^, were cultured in McCoy’s 5A Media (Corning, #10-050-CV) supplemented with 10% Fetal Bovine Serum (FBS) (Corning, #35-016-CV) and 1% Penicillin-Streptomycin (Gibco, #15240-062). Human mammary epithelial cells, MCF10A TP53^+/+^ and MCF10A TP53^-/-^ (Sigma-Aldrich) were cultured in 1:1 Dulbecco’s Modified Eagle Medium: Ham’s F-12 (Gibco, #11330-032) supplemented with 5% horse serum (Gibco, #16050-122), 20 ng/ml epidermal growth factor (Peprotech, #AF-100-15), 0.5 ng/ml hydrocortisone (Sigma, #H-0888), 100 ng/ml cholera toxin (Sigma, #C-8052), 10 µg/ml insulin (Sigma, #I-1882), and 1% Penicillin-Streptomycin (Gibco, #15240-062). The human near haploid cell line, HAP1 parental and HAP1 *ATF4^-^* cells (Horizon Genomics, HZGHC007380c010), were cultured in Iscove’s Modified Dulbecco’s Medium (Gibco, #12440-053) supplemented with 10% Fetal Bovine Serum (FBS) (Corning, #35-016-CV) and 1% Penicillin-Streptomycin (Gibco, #15240-062). All cell lines were cultured at 37°C and 5% CO_2_ in a water-jacketed incubator.

For cell line treatments, cells were cultured for times indicated in each experimental figure/legend with either 5 µM nutlin-3A (Millipore Sigma, #45-SML0580) to stabilize p53 activation, 100 µM etoposide (Thermo Scientific, #J63651.MC), 2 µM tunicamycin (Thermo Scientific, #J62217.MA) or 2 mM histidinol (Acros Organics, #AC228831000). All drugs were freshly resuspended in DMSO and DMSO-only controls were added at equal volumes to each drug treatment.

### Luciferase plasmid cloning and expression assays

Relevant plasmids and primers with cloning design are listed in Table S1. All cloning was done using NEBuilder (NEB, #E2621S). NanoLuciferase reporter plasmids were constructed using GADD45a_pHG plasmid as a backbone (kind gift from A. Fornace). pGB7 is the wild-type equivalent plasmid used throughout this study and includes *GADD45A* region chr1:67682954-67690203 (hg38) with translationally fused *nLuc*. pGB7 also engineered to have PacI, Esp3I and MunI sites to facilitate removal of the 5’ UTR and/or promoter region.

Luciferase assays were carried out using Nano-Glo® Dual-Luciferase® Reporter Assay System (Promega #1620) following manufacturer’s recommendations. NanoLuciferase or firefly luciferase values were normalized to constitutively expressed firefly luciferase (fLuc), or nanoLuciferase (nLuc) levels that were generated by co-transfected pGL4.53 or pNL1.1 (Promega, #E5011, #N1441) plasmids, respectively.

### dCas9-KRAB gRNA cloning

All primers used to generate gRNAs are listed in Table S1. To generate a transfer plasmid for each dCas9-KRAB genomic target region, forward and reverse primer pairs were annealed and then cloned into the BsmBI restriction site in the pLV hU6-sgRNA hUbC-dCas9-KRAB-T2a-Puro vector (gift from C. Gersbach, Addgene #71236). HCT116 cell lines constitutively expressing dCas9-KRAB and appropriate targeting gRNA were generated as described in (Thakore et al., 2015). All treatments are described Material and Methods above and in the figure legend.

### Lentivirus production, purification and transduction

Lentiviral particles were packaged using HEK293FT cells seeded at a density of 250,000 cells per well in 6-well culture plates. In brief, 1 μg of pLKO.1-Puro TRC plasmid containing either non-targeting control shRNA (5’-CAACAAGATGAAGAGCACCAA-3’) or ATF4-targeted shRNA (5’-GCCTAGGTCTCTTAGATGATT-3’) was combined with 1 μg of packaging plasmids psPAX2 and pMD2.G, mixed at a molar ratio of 2:1. The plasmid mix was diluted in jetPRIME buffer (Polyplus Transfection, 89129-924) and transfection reagent, following the manufacturer’s protocol. pMD2.G and psPAX2 were gifts from Didier Trono (Addgene plasmid # 12259, #12260). Lentiviral supernatants were collected at 24 and 48 h post-transfection, supplemented with 8 μg/ml Polybrene, filtered through 0.45-μm nitrocellulose filters, and stored at −80 °C. Cells were transduced with lentivirus and were selected for viral infection via addition of 2 μg/mL of puromycin for 72 hours.

## Quantitative Real Time PCR (RT-qPCR)

Total RNA was isolated (Quick RNA, Zymo, #R1055) with on-column DNase I digestion for 30 minutes. Single-stranded cDNA was generated (High Capacity cDNA Reverse Transcription Kit, ABI #4368814) and qPCR was performed using the relative standard curve method and iTaq Universal SYBR Green Supermix reagents (BioRad). All RT-qPCR primers are presented in Table S1.

### Western Blotting

Total protein was isolated using a custom RIPA buffer (50mM Tris-HCl[pH 7.4], 150mM NaCl, 1% NP–40, 0.5% sodium deoxycholate, 0.1% SDS) supplemented with protease/phosphatase inhibitors (Pierce, 78442). Protein concentration was measured using the BCA approach (Pierce, 23227), and equal protein concentrations were analyzed using the ProteinSimple® Wes platform with the 12–230 kDa Wes Separation Module containing 8 × 25 capillary cartridges per manufacturer’s instructions. Specific antibodies used were: anti-p53 (clone DO-1, BD Bioscience #554293), anti-ATF3 (Abcam, #AB207434), anti-ATF4 (Cell Signaling, #D4B8), anti-GADPDH (Cell Signaling, #5174S).

### CUT&RUN

1.5×10^6^ cells per CUT&RUN reaction and three biological replicates per condition were prepared for batch processing using the Epichyper CUTANA™ CUT&RUN protocol v1.9 and reagents (Epicypher, #14–1048). Briefly, cells bound to Concanavalin A beads were incubated overnight with 0.5 µg of anti-ATF4 antibody or non-specific rabbit IgG. DNA fragments were purified using phenol/chloroform extraction and 20 ng of purified DNA was used to construct an Illumina-compatible sequencing library (Liu, 2019) optimized for CUT&RUN-sized DNA fragments and NEBNext Ultra II DNA Library reagents (NEB #E7660). Library concentrations were quantified (NEBNext Library Quant Kit, E7630), pooled at equimolar concentrations, and sequenced on the Illumina NextSeq 2000 at the University at Albany Center for Functional Genomics.

### CUT&RUN and ChIP-seq Data Analysis

Raw paired-end sequencing reads for CUT&RUN were aligned to the hg38 human genome reference using hisat2 (Kim et al., 2019) with the following options (–X 700 –I 10 ––no-spliced-alignment). Regions of significant ATF4 enrichment (relative to IgG control signal) were identified using *macs2* (Zhang et al., 2008). ChIP-seq reads (Andrysik et al., 2017) for HCT116 input (GSM2296270), p53 ChIP-seq under DMSO-treatment (GSM2296271), and p53 ChIP-seq under nutlin-3A-treatment (GSM2296272) were downloaded from Gene Expression Omnibus and aligned to the hg38 human genome reference using hisat2.

BigWig files for visualization were produced via deepTools (Ramírez et al., 2016). Gene set enrichment for ATF4 CUT&RUN peaks was performed using ChIP-Enrich based on distance to the nearest Transcriptional Start Site (TSS) (Welch et al., 2014).

### RNA Sequencing

Cells were treated with either DMSO, nutlin-3A, etoposide, tunicamycin, or histidinol as described above in a six-well plate for 6h and total RNA was isolated (Quick RNA miniprep, Zymo, #R1055). PolyA+ RNA was purified using Dynabeads Oligo (dt)_25_ (Invitrogen, #61012) and fragmented at 94°C for 15 min. Fragmented RNA was used as the template for double-stranded cDNA production which was then used to construct an Illumina-compatible sequencing library (NEBNext Ultra II Directional RNA Library, NEB E7760). Libraries were then pooled for sequencing on an Illumina NextSeq 2000 at the University at Albany Center for Functional Genomics or on an Illumina Hiseq 2000 at Azenta/GeneWiz. Transcript abundance from the ENSEMBL hg38 genome assembly (v.104) was quantified using *kallisto* (quant –b 100) (Bray et al., 2016). Resulting transcript counts (TPM) were imported and processed via tximport (Soneson et al., 2015) and differential expression was quantified using DESeq2 (Love et al., 2014). Pathway enrichment analyses for differentially expressed genes were performed using enrichr (Chen et al., 2013; Kuleshov et al., 2016; Xie et al., 2021). Upstream regulator analysis on differentially-expressed genes was performed using the Causal Inference Engine (Farahmand et al., 2019) querying the TRRUST database (Han et al., 2018) using Fisher’s Exact Test settings and the STRINGdb (Szklarczyk et al., 2022) with the Quaternary Scoring statistic (Fakhry et al., 2016).

### STARRSeq enhancer mutagenesis screen

STARRSeq library preparation was done following a published protocol (Neumayr et al., 2019) with minor modifications as described here. To simplify STARRSeq library preparation, hSTARR-seq_ORI vector (Addgene #99296) (Muerdter et al., 2018) was modified by adding partial Illumina P5 and P7 adaptor sequence before AgeI and after SalI restriction sites, respectively, yielding plasmid pGB118. Mutagenesis library was constructed as a 250 nt of *GADD45A* intronic enhancer region (chr1:67686701-67686950) with a ‘N’ mixed base or a deletion at every position and 25 nt pGB118 matching overhangs on the 5’ and 3’ ends. The library was ordered as an oligo pool (oPool, IDT). oPool library was amplified for 10 cycles with primers SL1947 + SL1948 and cloned into pGB118 cut with AgeI and SalI using NEBuilder (NEB, #E2621S). 18 million HCT116 *TP53^+/+^* or *TP53^-/-^* cells were transfected with 10 µg STARRSeq mutagenesis plasmid library using JetPrime Transfection Reagent (Polyplus #101000046) following manufacturer’s recommendations. 5h after transfection, the media was replaced with fresh media supplemented with 0.001% DMSO, 5 µM nutlin-3A or 2 µM tunicamycin. RNA was extracted 6h post-treatment. Processed RNA and plasmid DNA libraries were sequenced on Illumina NextSeq 2000 instrument at the University at Albany Center for Functional Genomics as 2×150 paired-reads. R1 and R2 reads were first merged using bbmerge tool from BBMap (Bushnell et al., 2017). Merged reads were then collected and counted using a strict string pattern matching the expected library sequences. Relative expression was calculated as an RNA/DNA ratio.

## Results

### ATF3 is induced by the Integrated Stress Response (ISR) in a p53-independent manner

Activating Transcription Factor 3 (*ATF3)* is an immediate-early response gene in the cellular adaptive-response network. *ATF3* mRNA is upregulated in response to cellular stress including both DNA damage and endoplasmic reticulum (ER) stress (Hai et al., 1999; Hashimoto et al., 2002; Jiang et al., 2004; Hai, 2006; Lu et al., 2006). Recent reports suggest activation of the ISR leads to induction of p53 target genes, including *ATF3*, but the specific transcription factor requirements for this behavior have not been fully characterized. Thus, we investigated whether p53 was required for ATF3 induction under both DNA damage and ISR conditions. We confirmed *ATF3* mRNA expression is induced by both p53 and ISR pathways in HCT116 *TP53*+/+ (p53 WT) or *TP53*–/– (p53 null) colorectal carcinoma cells. We used two independent means to activate the p53 and ISR pathways. Etoposide activates p53 via induction of DNA double strand breaks (DSBs) (van Maanen et al., 1988; Shieh et al., 1997). Nutlin-3A specifically inhibits the negative p53 regulator, MDM2, leading to highly specific activation of p53 in a non-genotoxic manner (Vassilev et al., 2004). In vertebrates, the ISR is activated by ER stress, nutrient and heme deprivation, and viral infection, amongst others (Taniuchi et al., 2016). Therefore, we treated our HCT116 cell lines with tunicamycin, an inhibitor of N-linked glycosylation which induces ER stress via accumulation of unfolded proteins (Ding et al., 2007), or histidinol, which initiates the amino acid response (AAR) via depletion of histidine (Fu and Kilberg, 2013).

We first confirmed the specificity of these treatments via examination of p53 protein abundance and expression of canonical p53 and ISR target genes *CDKN1A/*p21 and the asparagine synthetase *ASNS*, respectively (el-Deiry et al., 1993; Szak et al., 2001; Siu et al., 2002). p53 protein (Fig. 1B) and *CDKN1A/*p21 mRNA (Fig. 1D) expression increased in response to both etoposide and nutlin-3A in a p53-dependent manner but not after ISR activation by tunicamycin or histidinol. Both ER stress and AA starvation led to *ASNS* induction in a p53-independent manner (Fig. 1E). Neither nutlin-3A nor etoposide treatments altered *ASNS* mRNA abundance in either genetic background, suggesting the ISR is not activated after DNA DSB induction. Taken together, these results suggest tunicamycin– and histidinol-mediated induction of the ISR in HCT116 cells does not require p53 activity (Fig. 1D, E). Although the induction of *ASNS* mRNA under ISR stimulating conditions relative to DMSO vehicle control was not dependent on p53, we note that the total abundance of *ASNS* mRNA is slightly reduced in HCT116 p53 null cell line (Fig. 1E).

**Figure 1.**
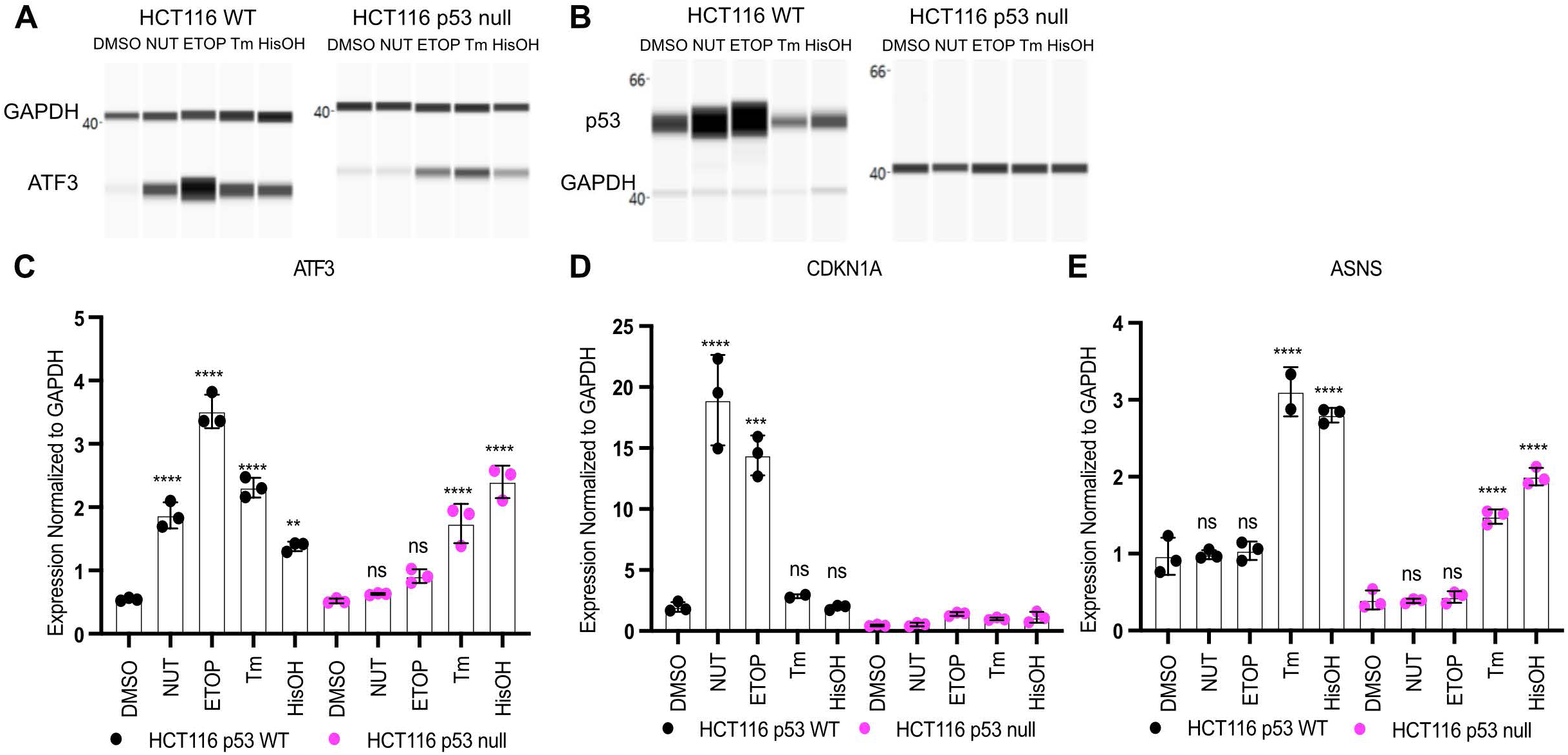
ATF3 is a p53 target gene that is activated via the ISR in a p53-independent manner. Western Blot analysis of A) ATF3 and B) p53 with GAPDH as a loading control in HCT116 p53 WT (left) and p53 null cells (right) following a 6h treatment with DMSO, 5 μM Nutlin-3A (NUT), 100 μM etoposide (ETOP), 2 μM tunicamycin (TM) or 2mM histidinol (HisOH). Gene expression analysis of the C) *ATF3* gene D) *CDKN1A* gene and E) *ASNS* gene in HCT116 p53 WT (black) and HCT116 p53 null (pink) cells in response to 6h treatment with stimuli. All statistical comparisons were computed using a one-way ANOVA test. *p<0.05, **p<0.01, ***p<0.001, ****p<0.0001.

*ATF3* mRNA and protein levels increased under both p53 and ISR stimulating treatments in HCT116 WT cells (Fig. 1A, C). p53 is required for induction of *ATF3* mRNA in response to nutlin-3A and etoposide, whereas ATF3 protein is still induced in response to etoposide in the absence of p53. Total ATF3 protein abundance is considerably lower in HCT116 p53 null cells relative to HCT116 WT suggesting that p53 activity is required for basal, but not induced, *ATF3* expression. Importantly, *ATF3* mRNA and protein induction is p53-independent under ISR stimulating conditions (Fig 1A, C). Coupled with the lack of p53 stabilization and *CDKN1A/p21* induction, these data suggest that upregulation of *ATF3* in response to ISR does not require p53.

### ATF4 and p53 independently regulate expression of ATF3

Our results suggest that both p53 and the ISR regulate *ATF3* transcription in a parallel, potentially redundant fashion. ATF4, a member of the basic region-leucine zipper (bZIP) TF superfamily, is one of the main transcriptional effectors of the ISR (Chen et al., 1996; Han et al., 2013). Prior work suggests ATF4 regulates expression of *ATF3* in other cellular contexts, therefore we tested whether ATF4 controls p53-independent induction of *ATF3* mRNA expression in response to ER stress and AA starvation (Pan et al., 2007; Kilberg et al., 2009; Fu and Kilberg, 2013). We first characterized the activity of *ATF4* in response to ISR-activating stimuli in our HCT116 p53 WT and p53 null cells to confirm ISR-dependent *ATF4* expression. As expected, *ATF4* mRNA and protein levels increase in response to both ER stress (via tunicamycin treatment) and AA starvation (via histidinol treatment), whereas *ATF4* mRNA and protein expression were unaffected in response to p53 stabilization (via nutlin-3A treatment) or DNA damage (via etoposide treatment) (Fig. 2A, D).

**Figure 2.**
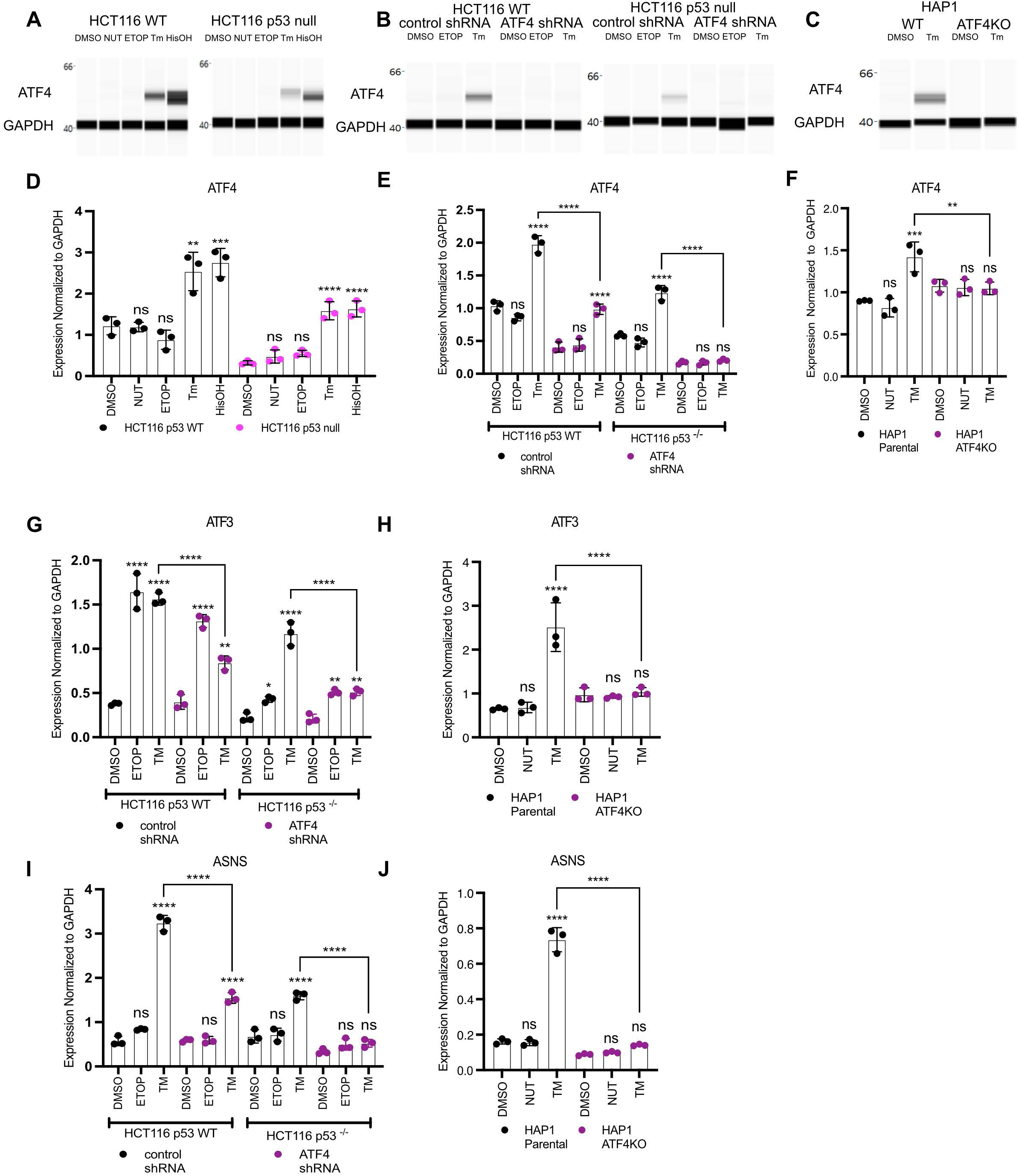
ATF4 and p53 independently regulate expression of *ATF3*. ATF4 protein (A-C) and mRNA (D-F) expression analysis by western blot and qRT-PCR, respectively, in HCT116 WT or p53 null cells (A, D), with shRNA constructs targeting control region (control shRNA) or *ATF4* (ATF4 shRNA) (B, E), or HAP1 WT or HAP1 ATF4KO cells (C, F). Gene expression analysis of *ATF3* (G, H) or *ASNS* (I, J) in HCT116 ATF4 shRNA cells (G, I) or HAP1 ATF4KO cells (H, J). Cells were harvested 6h post-treatment with DMSO, 10 μM nutlin-3A (NUT), 100 μM etoposide (ETOP), 2 μM tunicamycin (TM) or 2 mM histidinol (HisOH). All statistical comparisons were computed using a one-way ANOVA test. *p<0.05, **p<0.01, ***p<0.001, ****p<0.0001.

To determine the role of ATF4 in regulating *ATF3* induction downstream of ISR activation, we created HCT116 p53 WT and p53 null cells expressing either a non-targeting control shRNA or *ATF4*-targeted shRNA. *ATF4* mRNA and protein are substantially reduced in *ATF4* shRNA-expressing cells compared to non-targeting control shRNA (Fig. 2B, E), as was the ATF4 target *ASNS,* demonstrating the effectiveness of shRNA in ablating ATF4 (Fig. 2I). *ATF3* induction in response to etoposide was unaffected by *ATF4* depletion (Fig. 2G). Conversely, *ATF4* knockdown significantly reduced *ATF3* mRNA induction in response to ER stress, suggesting a direct role for ATF4 activity in mediating ISR-dependent *ATF3* expression (Fig. 2G). We extended our analysis to isogenic *ATF4*+ (HAP1 parental) and *ATF4*– (HAP1 ATF4KO) haploid leukemia cell lines. *ATF4* mRNA and protein levels were not induced in response to ER stress in HAP1 *ATF4-* cell line (Fig. 2C, F). Undetectable ATF4 protein in the *ATF4*– cell line results from a 2 bp insertion in exon 3 of *ATF4* that truncates the protein, including the DNA-binding domain. Lack of *ATF4* mRNA induction during ER stress in *ATF4-* cells suggests that ATF4 protein might be involved in *ATF4* autoregulation, however there is currently no evidence suggesting direct regulation (Dey et al., 2010). Inactivation of ATF4 led to a complete ablation of both *ASNS* and *ATF3* mRNA induction in response to ER stress, confirming *ATF3* induction in response to ISR is ATF4-dependent (Fig. 2H, J). These results demonstrate that ATF4 is the main effector of ISR-mediated *ATF3* induction and that p53 mediates *ATF3* transcription after DNA damage.

### ATF4 and p53 occupy distinct regulatory regions in the ATF3 gene locus

Our results demonstrate a genetic dependence for p53-mediated activation of *ATF3* transcription under DNA damage conditions, and a functionally distinct, ATF4-dependent pathway that regulates *ATF3* transcription during the ISR. To understand whether direct ATF4 binding to regulatory regions controls *ATF3*, we generated novel cleavage under targets & release under nuclease (CUT&RUN) (Skene and Henikoff, 2017) data for ATF4 in HCT116 cells treated with either DMSO (control), p53-activating (etoposide), or ISR-stimulating agents (tunicamycin or histidinol) using either an ATF4-specific antibody or a non-specific IgG control. We generated high-confidence, ISR-activated ATF4 binding events by considering called peaks from 5 out of 6 experiments from cells treated with either tunicamycin or histidinol (Fig. S1A-C). The rationale for this peak filtering strategy is to restrict further analysis of universal ISR-dependent ATF4 binding events, while also accounting for biological or technical variability in replicates. In support of this approach, we observe 7,723 ATF4 binding events shared across five out of six experimental conditions, with 5,093 (65%) existing across all observations. These 7,723 peaks were then examined for expected features of ATF4 binding, including specificity during ISR stimulation and enrichment of predicted ATF4 DNA binding motifs.

Known motif enrichment analysis revealed the ATF4 motif as enriched in high-confidence peaks, followed by enrichment of the known heterodimer partner, C/EBP homologous protein (CHOP) (Fig. 3A). Similar enrichment of a motif most closely matching the known ATF4 motif was observed using *de novo* enrichment strategies (Fig. 3B). Enrichment of CUT&RUN tags was highly specific for tunicamycin and histidinol treatment conditions compared to either vehicle DMSO or etoposide conditions (Fig. 3C). To provide context into the functional relevance of ATF4 binding, our CUT&RUN analysis reveals a significant, ISR-specific enrichment of ATF4 at the *ASNS* promoter (Fig. S1D), confirming previous reports that AA starvation-induced *ASNS* expression is mediated via ATF4 interaction with the promoter (Chen et al., 2004), We also assessed whether the ATF4 high-confidence binding was associated with genes in the ISR or other known pathways. We used ChIP-Enrich to perform gene pathway enrichment analyses on genes nearest to either our ATF4 binding events or previously identified p53 binding events from nutlin-3A-treated HCT116 cells (Welch et al., 2014; Andrysik et al., 2017). Genes in the Unfolded Protein Response and aminoacyl-tRNA biosynthesis pathways are highly enriched gene sets associated with ATF4 binding when querying the mSigDB, KEGG, and REACTOME databases (Fig. 3E-G), in support of our ATF4 CUT&RUN data representing functional binding events. p53 binding events are associated with genes involved in apoptosis, DNA repair, and the response to stress, as expected (Fig. 3E-G). Interestingly, these analyses revealed both ATF4 and p53 binding events are significantly associated with genes in the canonical p53 pathway, including *ATF3* (Fig. 3E-F). Taken together, these data suggest that our high-confidence, ISR-dependent ATF4 peaks likely represent true ATF4 genomic binding events and further suggest a link between the ATF4-dependent ISR and the p53 network.

**Figure 3.**
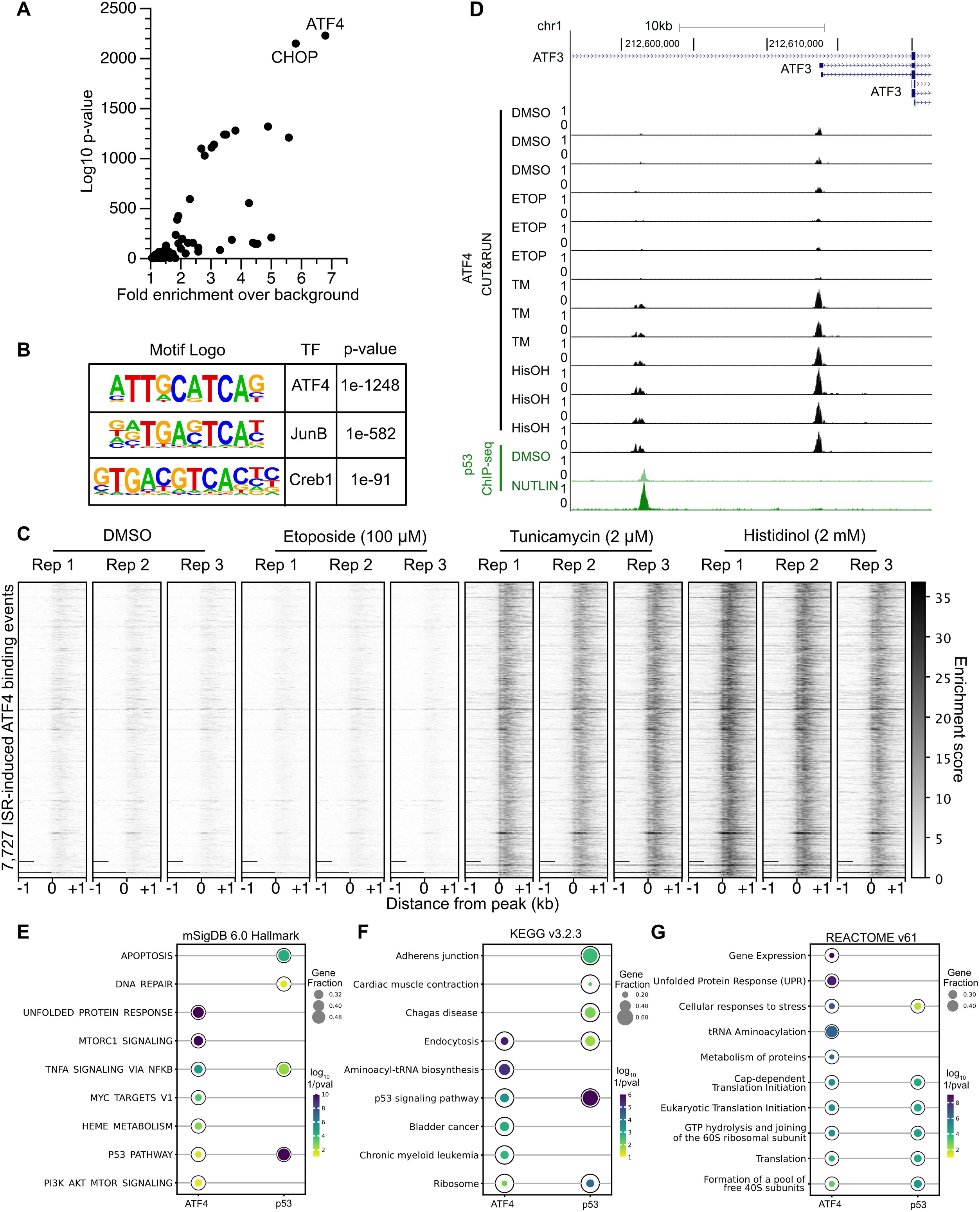
ATF4 and p53 occupy distinct regulatory regions in the *ATF3* gene locus. A) Known motif enrichment analysis of the high-confidence peak set reveals the predicted ATF4 motif as the most highly enriched motif within this dataset. B) *de novo* motif analysis of high-confidence peak set shows enrichment of ATF4 motif. C) Enrichment of CUT&RUN sequencing tags for the 7,723 high-confidence ATF4 peaks after 6h drug treatments as indicated from –1 kb and +1 kb from peak center. D) Genome browser view of the *ATF3* gene locus displaying ATF4 CUT&RUN data (black) and p53 ChIP-Seq data (green) (scaled to 1 as the maximum value for ATF4 or p53) following 6 h treatment with various stress stimuli: DMSO (vehicle control), 5 μM nutlin-3A (NUTLIN), 100 μM etoposide (ETOP), 2 μM tunicamycin (TM), or 2mM histidinol (HisOH). E-G) The top 5 most enriched results from *chiprenrich* for ATF4 CUT&RUN high-confidence peaks (from this manuscript) or nutlin-induced p53 ChIP-seq peaks (from (Andrysik et al., 2017)) for E) the mSigDB v6.0 Hallmark (Liberzon et al., 2015), F) KEGG v3.2.3 (Kanehisa et al., 2023), or G) REACTOME v61 (Gillespie et al., 2022) gene sets. *P*-values are log_10_ (1/ Bonferroni-corrected P-value). Full data tables for enrichment results can be found in Table S5.

Given the genetic evidence that *ATF3* is regulated by both p53 and ATF4 and our genome-wide ATF4 binding experiments, we next surveyed the *ATF3* gene locus to identify putative ATF4 and p53 binding events that might directly regulate *ATF3* transcription. We observe ATF4 binding within a previously identified promoter regulating ISR-dependent *ATF3* transcription (Fu and Kilberg, 2013) in response to both ER stress and amino acid deprivation, but not during the DDR or in DMSO control conditions (Fig. 3D). p53, on the other hand, binds to a DNAse Hypersensitive Site (DHS) approximately 13 kb upstream (p53-bound DHS) from the ATF4-bound *ATF3* promoter. ATF4 also occupies a spatially distinct DHS 15kb upstream (ATF4-bound DHS) from the *ATF3* TSS in an ISR-dependent fashion. These novel genomic binding datasets for ATF4 further suggest a direct role for ATF4 in regulation of ISR-dependent *ATF3* and confirm the ATF4-independence of *ATF3* transcription downstream of DNA damage.

### Analysis of the regulatory elements controlling stress dependent ATF3 expression

Biochemical analyses and genetic loss-of-function experiments confirm p53 and ATF4 independently regulate expression of *ATF3* (Figs. 1-3). To determine if the distinct p53 or ATF4-bound DHS contribute directly to transcription of *ATF3* in response to stress, we tested their ability to activate transcription of a luciferase reporter gene via a minimal promoter (minP). The upstream ATF4-bound DHS does not act as an enhancer, as there was no significant difference in transcription compared to the negative control (minP only), under any conditions and cellular contexts tested (Fig. 4A). The p53-bound DHS drove substantial transcriptional activity under basal conditions with a significant increase in activity upon nutlin-3A treatment. Basal and p53-induced activity were significantly reduced when the putative p53 binding motif was mutated or when assayed in HCT116 p53 null cell lines. Loss of p53 does not completely ablate basal transcription, suggesting other transcription factors likely contribute to unstimulated *ATF3* activity. However, no additional transcriptional activation by the p53-bound DHS was observed in response to tunicamycin (Fig. 4A). Taken together, these data suggest that the p53-bound DHS likely regulates *ATF3* transcription under basal and conditions that stabilize p53, and that the adjacent ATF4-bound DHS likely does not facilitate ISR-mediated *ATF3* transcription.

**Figure 4.**
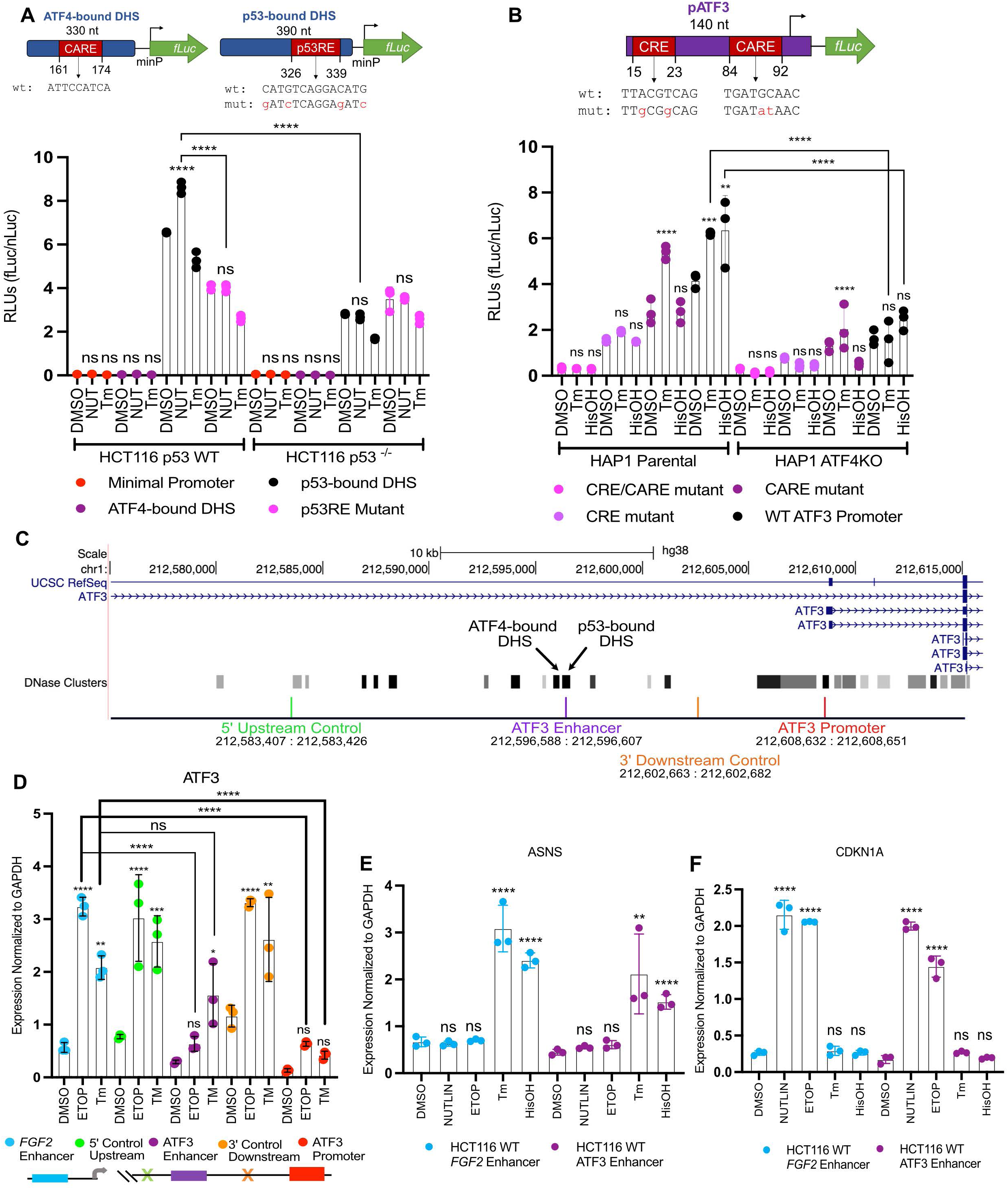
*ATF3* induction by the ISR does not require the upstream enhancer element bound by p53. A) Normalized luciferase values driven by the upstream *ATF3* DNase Hypersensitivity sites (DHS): ATF4-bound DHS, p53-bound DHS, p53RE Mutant, and the minimal promoter (negative control), in response to 16h treatment with DMSO, 5 μM nutlin-3A (NUT) or 2 μM tunicamycin (Tm) in HCT116 p53 WT and p53 null cells. B) Normalized luciferase values driven by the (–104/+36) *ATF3* promoter sequence (WT *ATF3* promoter) and constructs containing mutations in specific ATF4 response elements: CARE, CRE, CARE/CRE, in response to 16 h treatment with DMSO, 2 μM tunicamycin (Tm), or 2mM histidinol (HisOH) in HAP1 parental and ATF4KO cells. Luciferase reporters with relevant motif positions and sequences are illustrated above the corresponding bar chart (A-B), genomic locations of the *ATF3* promoter and DHS are reported in Table S1. C) Genome browser view of the *ATF3* gene locus displaying the location of dCas9-KRAB gRNA targets and the genomic coordinates spanning these targets relevant to panel D. D) RT-qPCR analysis of the *ATF3* gene in response to 6 h treatment with DMSO, 100 μM etoposide (ETOP) or 2 μM tunicamycin (TM) in HCT116 p53 WT cells where dCas9-KRAB is targeting regions at off-target sites at a control FGF2 enhancer (blue) or intergenic region (orange and green), the p53-bound *ATF3* enhancer element (purple) or *ATF3* promoter (red) for transcriptional repression. RT-qPCR analysis of the E) *ASNS* gene, and F) *CDKN1A*/*p21* gene, following a 6h treatment with various stress stimuli. Statistical comparisons for nascent expression levels were computed using a one-way ANOVA test. *p<0.05, **p<0.01, ***p<0.001, ****p<0.0001.

Prior work suggests ATF4 regulates *ATF3* via interaction with two canonical ATF/CREB family motifs within the *ATF3* promoter in hepatocarcinoma (HepG2) cells (Fu and Kilberg, 2013). This regulation depends on both a CRE site (nt −90/−82, TTACGTCAG) and a CARE site (nt −21/−13, TGATGXAAX) within the *ATF3* gene promoter (−104/+36) (Weidenfeld-Baranboim et al., 2009; Hai et al., 2010). We thus assessed whether ATF4 regulates *ATF3* via interaction with these elements, as ISR-induced ATF4 binding to an upstream DHS had no effect on *ATF3* transcription (Fig. 4A). We thus tested activity of luciferase reporters driven by the −104/+36 promoter fragment of the *ATF3* gene – containing both the CARE and CRE sequences (WT ATF3 promoter), as well as constructs containing mutations in one (CARE and CRE) or both (CARE/CRE) of these sites, in control and ATF4-deficient HAP1 cell lines. Consistent with prior reports, both ER stress (tunicamycin) and AA starvation (histidinol) treatments led to increased transcription driven by the WT *ATF3* promoter in an ATF4-dependent manner (Fig. 4B). Mutation of the CARE and CRE sites, both capable of supporting ATF4 binding, showed significantly and substantially reduced ability to drive both basal and ISR-induced transcription. Our data are consistent with prior observations that the CARE site is required for amino acid starvation-induced activity, but is dispensable for ER stress-induced transcription (Pan et al., 2007; Fu and Kilberg, 2013). Similar to our results for the upstream p53-bound DHS, these data suggest that multiple transcription factors in addition to p53 and ATF4 are likely involved in the basal and stress-dependent regulation of *ATF3*, consistent with models whereby multiple transcription factors work in a context-dependent and combinatorial manner to drive transcription (Smith et al., 2013; Chaudhari and Cohen, 2018; Zeitlinger, 2020; Choi et al., 2021; Kim et al., 2021). Our data using genetic depletion strategies, biochemical binding assays, and testing putative *cis-* regulatory element activity using reporter assays provide evidence for a model whereby *ATF3* transcription in response to DNA damage and the ISR occurs through distinct transcription factors binding to distinct regulatory elements.

### ISR-mediated induction of ATF3 does not require the upstream enhancer bound by p53

ATF4 and p53 likely independently regulate expression of *ATF3* by occupying distinct putative regulatory regions in a stress-dependent manner (Figs. 2G, 3D, 4A-B). To further characterize the mutual independence of p53 and ATF4 and to demonstrate whether these binding events control *ATF3* transcription *in vivo*, we utilized a CRISPR interference (CRISPRi) system to block effector protein binding at these elements (Gilbert et al., 2013; Qi et al., 2013). We chose the dCas9-KRAB CRISPRi system which fuses a catalytically inactive *Streptococcus pyogenes* Cas9 with a KRAB transcriptional repressor domain (Margolin et al., 1994). dCas9-KRAB targeting to *cis*-regulatory elements is an effective strategy for blocking effector protein binding and inhibiting regulatory elements and linked gene expression (Thakore et al., 2015; Yeo et al., 2018; Catizone et al., 2020). We first targeted dCas9-KRAB to the *ATF3* promoter as proof of principle, since repression of a gene promoter should inhibit transcription initiated from that element. Targeting of dCas9-KRAB to the *ATF3* promoter, approximately 100 bp upstream of the TSS significantly reduced *ATF3* mRNA levels (Fig. 4D). *ATF3* mRNA levels were unaffected when targeting dCas9-KRAB to the off-target p53-bound *FGF2* enhancer (chr4:122,798,358, *hg38*) or a region 13kb upstream (5’) of the putative *ATF3* enhancer (Fig. 4D). A similar lack of repression was observed when dCas9-KRAB is targeted to a region equidistant (6kb) from both the putative *ATF3* enhancer and promoter (3’ downstream relative to the enhancer) (Fig. 4D). This repression when targeting the promoter was observed under basal (DMSO), DNA damage, and ER stress conditions, demonstrating the effectiveness of the CRISPRi system. Targeting dCas9-KRAB to the p53-bound enhancer significantly reduced *ATF3* mRNA levels in response to basal and etoposide-treated conditions when compared to all non-targeting controls (Fig. 4D). Targeting of dCas9-KRAB to any of the three control locations did not significantly alter either basal or DNA damage-induced *ATF3* expression (Fig. 4D). Canonical ISR and p53 gene targets *ASNS* and *CDKN1A/*p21 were unaffected by targeting dCas9-KRAB to the p53-bound enhancer or control regions (Fig. 4E-F). Induction of *ATF3* mRNA in response to tunicamycin-induced ER stress was not affected when targeting the p53-bound enhancer (Fig. 4D). These results indicate the p53-bound enhancer is important for both basal and p53-mediated activation of *ATF3*, but is not directly required for ER stress-mediated induction of *ATF3*.

### Global transcriptome analysis identifies common gene regulatory targets of the p53 and Integrated Stress Response

Our data demonstrate that the DNA damage response (DDR, via p53) and the Integrated Stress Response (ISR, via ATF4) independently activate transcription of *ATF3* through at least two gene regulatory elements. We sought to determine if this parallel stress-dependent gene activation by the p53 and ISR pathways may be more widespread. We thus performed polyA+ RNA-seq on HCT116 p53 WT and p53 null cells after 6 hours of treatment with p53 or ISR-activating stimuli: DMSO (vehicle), 5 μM nutlin-3A,100 μM etoposide, 2 μM tunicamycin, or 2 mM histidinol. Three biological replicates were analyzed for each treatment condition via transcript counting (*kallisto*, 100 bootstraps) (Bray et al., 2016) and differential gene expression analysis (*deseq2)* (Love et al., 2014). We confirmed that each treatment elicited an expected transcriptional response using gene ontology analysis of significantly upregulated (Fig. S2A-D) or downregulated (Fig. S2E-H) genes when compared to DMSO control (Chen et al., 2013; Kim et al., 2019; Xie et al., 2021). Treatment with either nutlin-3A or etoposide led to significant upregulation of genes (184 and 1,394, respectively) consistent with a functional p53 response, including those related to the cellular response to DNA damage (Fig. S2A-B). Treatment with tunicamycin led to upregulation of 2,138 genes consistent with ER stress and transcriptional regulation (Fig. S2C) and downregulation of 2,131 genes involved in translation and ribosome biogenesis (Fig. S2G). Similar ontology groups were enriched in differentially regulated genes after treatment with histidinol (Fig. S2D,H), although we note treatment specific enrichment of ER stress-associated genes after tunicamycin treatment and metabolic regulation after histidinol addition. Nearly 50% of the genes upregulated in response to either tunicamycin or histidinol have an ATF4 binding event within 10kB of their transcriptional start site (Fig. 5B), further supporting that these genes are likely regulated via ATF4 and the ISR. Taken together, these broad analyses of gene regulation via standard gene ontology methods demonstrate that each chemical treatment recapitulates expected cellular responses to specific cell stress conditions.

**Figure 5.**
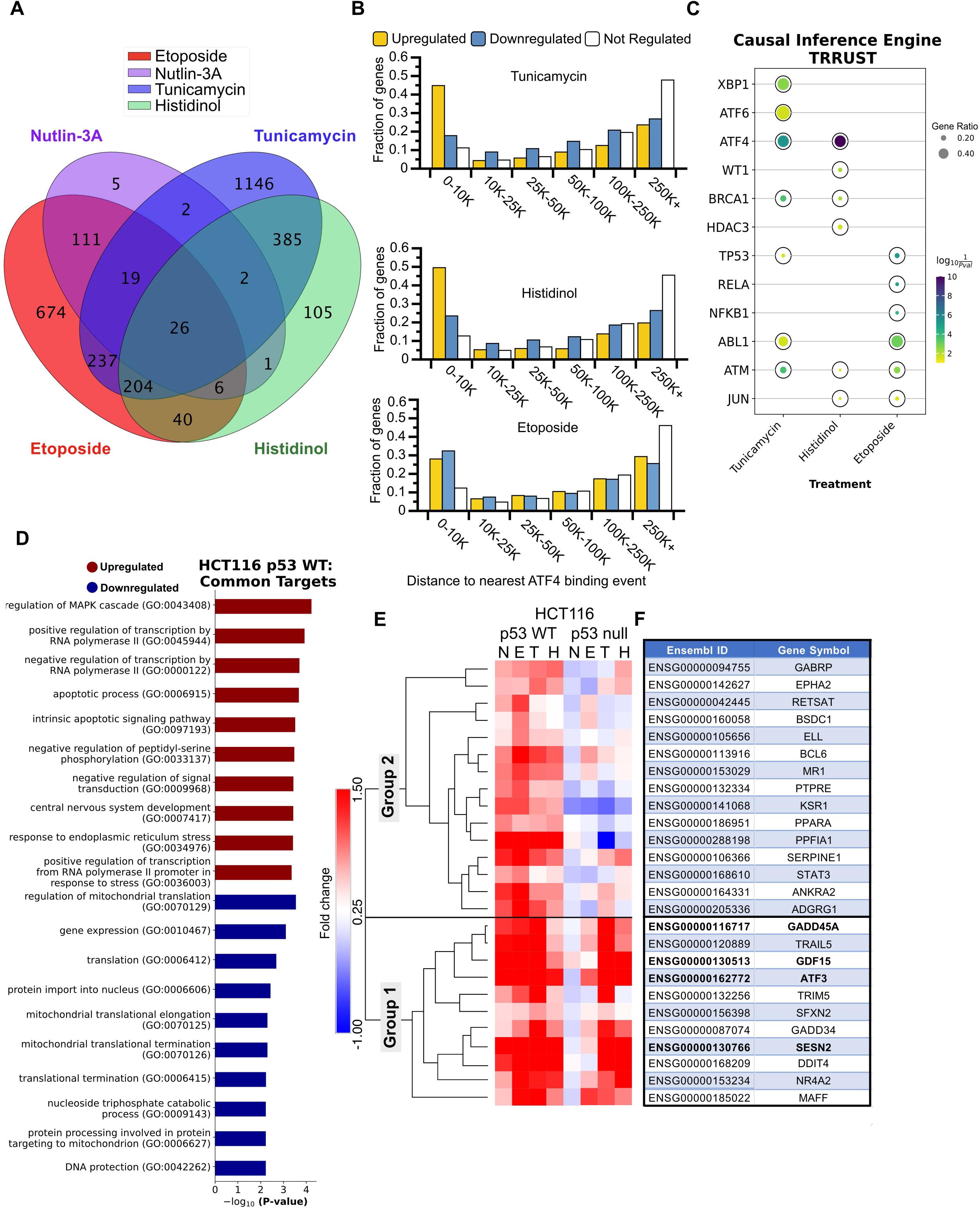
Global transcriptome analysis identifies common gene regulatory targets of the p53 GRN and the ISR. A) Intersection of genes upregulated (any fold-change relative to DMSO, Bonferroni-adjusted P-value <0.05) in HCT116 p53 WT cells treated with 5 μM nutlin-3A, 100 μM etoposide, 2 μM tunicamycin, and 2 μM histidinol, when compared to vehicle control (DMSO) for 6h. B) Bar plots representing a fraction of genes identified from the RNA-seq experiment as upregulated (yellow), downregulated (blue), or not regulated (white) in response to various stimuli that have a CUT&RUN-defined ATF4 binding event within a binned distance indicated on the *x*-axis. C) Causal Inference Engine predictions of the top 5 putative upstream regulators of genes from the TRRUST database induced by tunicamycin, histidinol, or etoposide treatment (as determined using the Fisher’s exact test for significance) (Han et al., 2018; Farahmand et al., 2019). Complete Causal Inference Engine results for the STRING database for both WT and HCT116 p53 null cells can be found in Fig. S3 and Table S4 (Szklarczyk et al., 2022). D) Gene ontology analysis of the genes commonly upregulated and downregulated (any fold-change relative to DMSO, Bonferroni-adjusted P-value <0.05), in response to various stress stimuli. E) Heatmap and hierarchical clustering results (one minus Pearson, average linkage) displaying fold change values for the 26 common targets identified in panel A. F) Table displaying the gene symbols for the 26 common target Ensembl gene IDs identified in panel A and matching row names in panel E. Genes validated in Fig. 6 by RT-qPCR are depicted in bold font.

We next wanted to identify additional genes that might be co-regulated by ATF4 and p53 and examine potential upstream regulators for both gene networks. We first used the Causal Inference Engine approach to identify potential upstream regulators of the differentially expressed genes identified in our RNA-seq analysis from the TRRUST network database (Han et al., 2018; Farahmand et al., 2019). ATF4 is predicted to be a significant regulator for both tunicamycin and histidinol-induced genes, but not etoposide in both WT and p53 null cell lines (Fig. 5C, S3A). ATF6 and XBP1, other transcription factors activated by ER stress, are also putative regulators of tunicamycin-induced genes, in agreement with their expected role in mediating ER stress (Pakos-Zebrucka et al., 2016). Interestingly, this approach also predicts p53 is a likely regulator of ISR-mediated gene expression in both WT and p53 null cells and across both the TRRUST and STRING networks (Fig. 5C, S3A-C). These data, combined with our extensive genetic and biochemical analyses of *ATF3* regulation, suggest additional genes may be co-regulated by these two core cellular stress responses via similar mechanisms to *ATF3*.

We therefore established parameters to define genes that can be independently activated by p53 and the ISR. We reasoned that these genes would be i) significantly upregulated in response to both p53– and ISR-activating stimuli in HCT116 p53 WT cells relative to DMSO, ii) p53-dependent in response to nutlin-3A treatment, a stimulus that specifically activates and stabilizes p53 (Fig. S2I), and iii) significantly upregulated in the absence of p53 in response to ISR-activating treatments, tunicamycin and histidinol. The inclusion of nutlin-3A-mediated regulation limits our gene set to those genes regulated by p53 specifically, as opposed to via DNA damage-dependent, but p53-independent mechanisms, as we have previously observed (Catizone et al., 2020). Ultimately, these criteria yielded 26 genes upregulated in response to p53 activation, ER stress, and AA starvation (Fig. 5A). As expected, *ATF3* was in this gene set upregulated in response to all four treatment conditions relative to DMSO, providing support for selection criteria and the quality of the data set (Fig. 5E-F). Gene ontology analysis of these 26 genes suggests shared involvement in apoptotic signaling pathways and transcription by RNA polymerase II. We also observed regulation of the mitogen-activated protein kinase (MAPK) signaling cascade, which integrates and amplify signals from diverse stimuli (Zhang and Dong, 2007; Chavel et al., 2010) (Fig. 5D). Pathway analysis on shared downregulated genes reveal common function in protein synthesis, potentially reflecting a switch from an anabolic to catabolic state, consistent with prior reports of broad translational control by these pathways (Fig. 5D) (Loayza-Puch et al., 2013; Zaccara et al., 2014; Andrysik et al., 2017; Tameire et al., 2019; Tian et al., 2021).

We next analyzed the behavior of commonly upregulated genes in HCT116 p53 null cells to determine whether induction is truly p53-dependent or independent in response to ISR-inducing stimuli. Heirarchical clustering (one minus Pearson correlation with complete linkage) revealed two distinct groups of these genes (Group 1 and Group 2). 11 of the 26 commonly upregulated genes behave similarly to *ATF3* (Figs. 5E-F) and cluster together in Group 1. The remaining genes (Group 2) show some dependence on p53 for full activation downstream of the ISR. p53 protein levels are not stabilized in response to ISR-activating stimuli (Fig. 1B) and we observe no evidence that canonical p53 targets like *CDKN1A*/p21 respond to ISR-activating stimuli (Fig. 1D), suggesting that this partial dependence on p53 is likely due to indirect activity of p53 in regulating other activators within the ISR. Consistent with this, we note diminished ATF4 protein in the absence of p53 in basal and ISR-induced conditions (Fig. 2A). While additional work is required to determine the causal relationship, if any, between loss of p53, reduced ATF4 protein abundance, and the effect on ISR-dependent gene expression, our data suggest that p53 and ATF4-dependent transcriptional control mechanisms are likely to be functionally independent.

Lastly, we validated the p53 and ATF4-dependence of a set of common genes using a battery of cell lines and genotypes. We examined p53-dependence for three candidate genes regulated similarly to *ATF3*: *GADD45A* (Zhan et al., 1994; Ebert et al., 2019)*, SESN2* (Budanov and Karin, 2008; Garaeva et al., 2016), and *GDF15* (Osada et al., 2007; Li et al., 2021). We tested the behavior of these three genes in response to etoposide or tunicamycin treatment in p53 WT and p53 null HCT116 (Fig. 6A-D) and MCF10A mammary epithelial lines (Fig. 6E-H). Each gene was induced by both treatments and etoposide-mediated induction required an intact p53 response. Conversely, p53 was not required for induction in response to tunicamycin, consistent with the parallel nature of the p53 and ATF4-dependent transcriptional networks. These data in MCF10A cell lines also demonstrate that this behavior is not limited to colon carcinoma cell lines. We then assessed ATF4-dependence of these genes in either HAP1 parental or *ATF4-* cell lines and found that *ATF4* activity is required for tunicamycin induction (Fig. 6I-L). Nutlin-3A treatment fails to induce expression of *ATF3, GADD45A, SESN2,* and *GDF15* likely due to the loss-of-function *TP53* S215G allele (Moder et al., 2017) in these cell lines. The absence of p53 and the presence of a functional ISR further suggest the functional independence of these pathways. Our data thus identify a set of “dual response” genes independently regulated by p53 or ATF4 in response to specific stress conditions.

**Figure 6.**
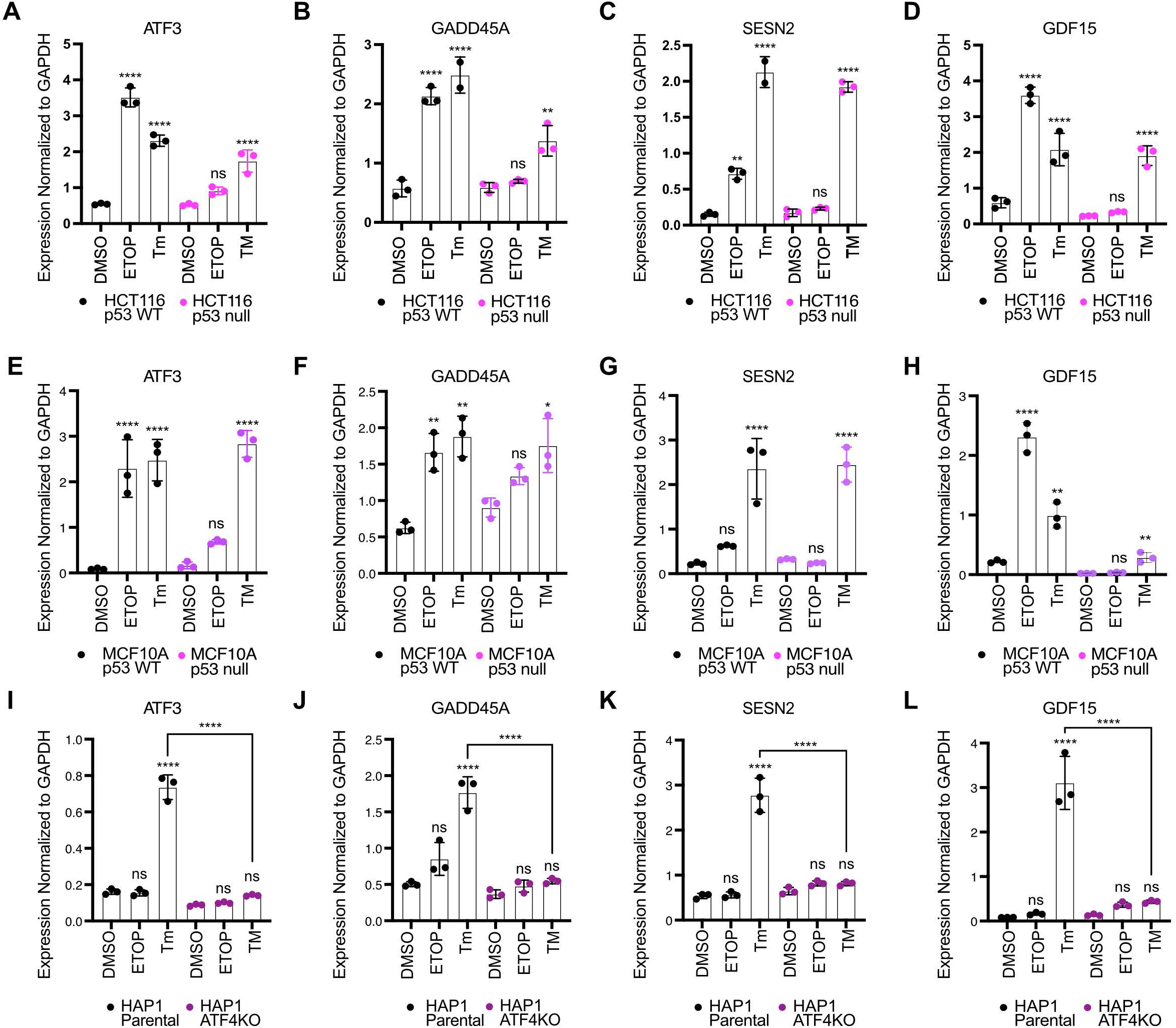
Parallel stress-dependent networks converge at activation of a common set of target genes. RT-qPCR analysis of the *ATF3, GADD45A*, *SESN2,* and *GDF15* gene in A-D) HCT116 p53 WT and p53 null cells, E-H) MCF10A p53 WT and p53 null cells, and I-L) HAP1 parental and ATF4KO cells, following a 6h treatment with DMSO, 100 μM etoposide (ETOP), or 2 μM Tunicamycin (Tm). All statistical comparisons were computed using a one-way ANOVA test. *p<0.05, **p<0.01, ***p<0.001, ****p<0.0001.

### GADD45A gene-derived reporter system for assessing DDR and ISR enhancers

Our data demonstrate p53 activates *ATF3* transcription through a distal enhancer, whereas we confirmed observations that ATF4 regulates transcription via the *ATF3* proximal promoter. We sought to determine whether any additional “dual response” genes have *cis*-regulatory strategies like *ATF3*. We focused on *GADD45A* which was reported to contain a p53-bound enhancer within the 3^rd^ intron (Zhan et al., 1998; Daino et al., 2006). Our CUT&RUN analysis suggests ATF4 also binds to this putative enhancer (Fig. 7A). We thus tested whether p53 and ATF4 binding to this element controls *GADD45A* transcription under p53 or ISR activating conditions, respectively, utilizing a newly constructed luciferase reporter system (Fig. 7B). We reasoned that (1) *GADD45A* is relatively small (∼3 kb) and is convenient for plasmid-based genetic manipulations, and that (2) this reporter might be more relevant for stress-dependent promoter-enhancer interaction studies as a near native genetic context, including enhancer:promoter distance and location, is maintained. We tested four versions of the “native” *GADD45A*-*nLuc* system, creating transcriptional or translational fusion constructs either with or without a degron tag (hPEST) (Fig. S4). Luciferase activity was detectable in all four versions under basal conditions, and constructs lacking the degron tag were inducible by p53 (nutlin-3A) and ISR stimulating (tunicamycin) conditions (Fig. S4B), consistent with our results measuring native GADD45A mRNA expression (Fig. 6B). All subsequent experiments utilize the *GADD45A*-*nLuc* translational fusion lacking the degron tag due to the highest signal-to-noise ratio in our initial tests (Fig. S4B).

**Figure 7.**
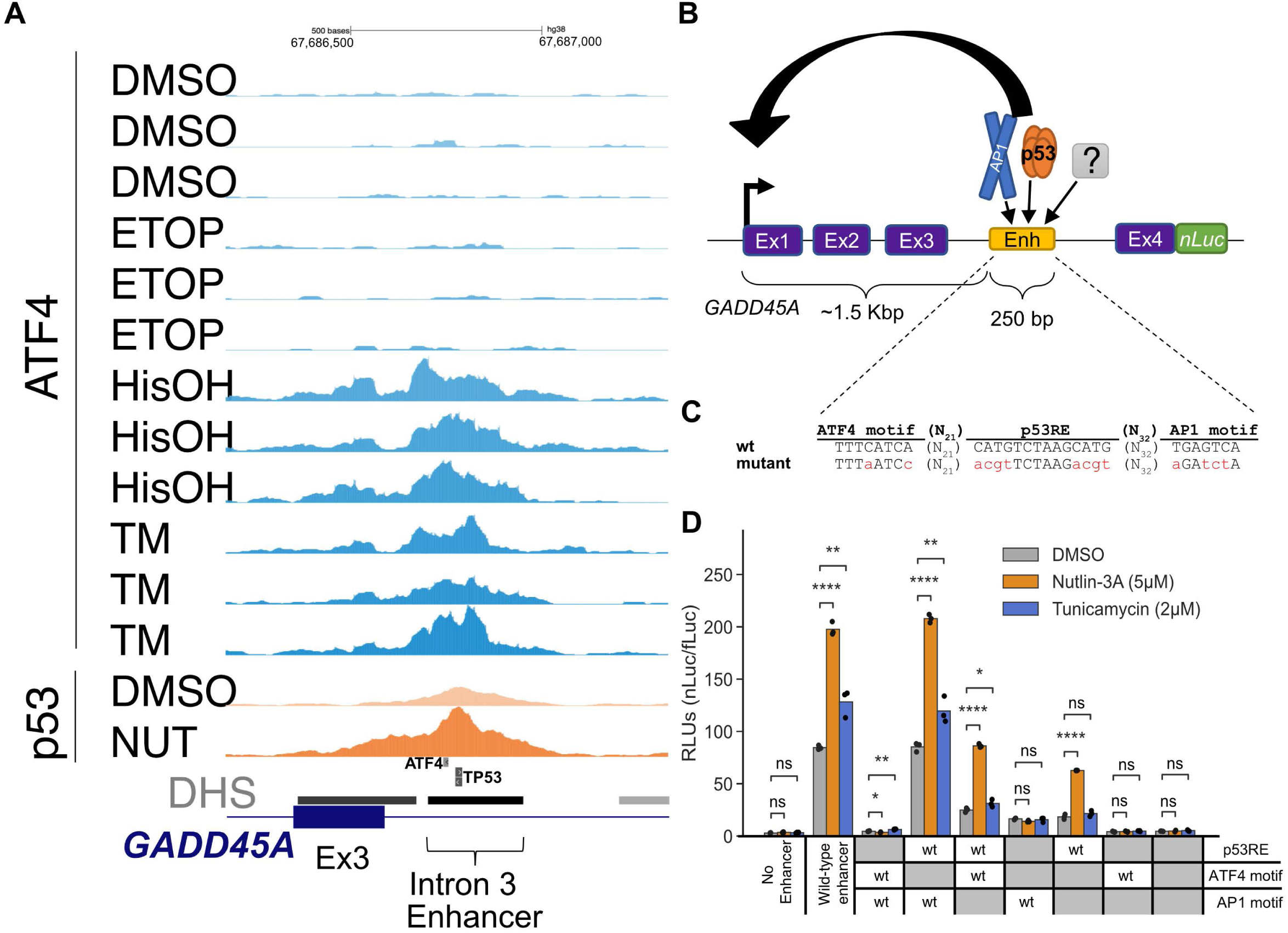
*GADD45A* as a reporter system to study DDR– and ISR-dependent enhancers. A) Genome browser view with *GADD45A* locus displaying ATF4 CUT&RUN and p53 ChIP-Seq data (Andrysik et al., 2017) in HCT116 WT cell line following 6h treatment with DMSO, 2 mM histidinol (HisOH), 2 μM Tunicamycin (TM), and 5 µM nutlin-3A (NUT). Putative p53RE, ATF4 motif and GADD45A intron 3 enhancer locations are indicated on the bottom, DNaseI hypersensitive sites (DHS) are marked in grey/black. B) Schematic representation of the *GADD45A-nLuc* reporter construct with relevant enhancer sequence motifs highlighted in panel C. D) Normalized luciferase expression values using *GADD45A-nLuc* reporter transfected into HCT116 wt cell line and 16h treatment with DMSO, nutlin-3A and tunicamycin as indicated in the legend. Reporter constructs included wild type, a negative control with 250 bp enhancer deletion (‘No Enhancer’) and various ATF4, AP1 and p53RE motif mutations alone or in combination as indicated in the table below (‘wt’ or ‘mutant’ in grey). Specific mutations in motifs are indicated in panel C. Statistical comparisons were generated using one-way ANOVA: * p<0.05, ** p<0.01, *** p< 0.001, **** p<0.0001.

To confirm the putative *cis*-element in the 3^rd^ intron is essential for *GADD45A-nLuc* reporter activity, we characterized luciferase activity in response to mutation or deletion of nucleotides predicted to be critical for binding of p53 and/or ATF4. The intronic enhancer has conserved p53RE and AP1 motifs that were previously reported to be required for ionizing radiation induced *GADD45A* expression (Chin et al., 1997; Daino et al., 2006; Smeenk et al., 2008). We additionally identified one canonical ATF4 motif (TGATGAAA, minus strand, Fig. 7C) using the JASPAR database (Castro-Mondragon et al., 2022). ATF/AP1 transcription family motifs are highly similar (Bejjani et al., 2019) (Fig. 7C, S5) and ATF4 can form heterodimers with other AP1 family members (Hai and Curran, 1991; Podust et al., 2001; Mann et al., 2013), thus we included mutants of both motifs to identify the true ISR response element in the *GADD45A* enhancer. The wild-type *GADD45A*-*nLuc* reporter system responds to the p53 and ISR pathways like the native *GADD45A* gene (Fig. 7D). An intact p53RE motif was required for maximal enhancer-driven transcription under basal conditions and consistent with prior work (Daino et al., 2006). Disrupting this motif (Fig. 7C) resulted in low expression levels like deletion of the entire 250bp DHS region suggesting that p53 is a major regulator of basal *GADD45A* transcription. Nutlin-3A-mediated transcription was completely dependent on the p53RE motif, whereas ATF4 and AP1 motifs were dispensable (Fig. 7D). Disrupting the predicted ATF4 motif had no effect on basal or tunicamycin-induced expression whereas disrupting the AP1 motif decreased basal activity and tunicamycin-mediated induction (Fig. 7D), suggesting that ISR pathway is regulated, at least partially, through the AP1 motif. Both the p53 and AP1 motifs appear to play key roles in the basal expression of *GADD45A,* but the motifs appear to be functionally distinct in response to p53 and ISR-stimulating conditions. We also included combinatorial motif mutants to investigate whether transcription factors binding to these motifs are activating independently. If the predicted ATF4 motif was not functional, one would expect that the double inactivation of p53RE and ATF4 motif would behave as p53RE mutant and would still be inducible by tunicamycin. In contrast, we observed lack of tunicamycin-mediated induction when the reporter had combined p53RE and ATF4 motif mutations. In fact, all constructs that included ATF4 and/or AP1 mutations in different combinations were not inducible by tunicamycin (Fig. 7D). It is possible that the ATF4 motif is not required under normal conditions but could act redundantly in the absence of the preferred AP1 motif.

### Nucleotide-level characterization of the GADD45A enhancer reveals critical regulatory sequences

Enhancers are generally regulated by multiple transcription factors (TFs) acting positively or negatively depending on context (Kim and Wysocka, 2023). p53 binding primarily positively regulates enhancer activity, although the extent to which additional transcription factors are required for p53-dependent *trans-* activation remains an open question (Verfaillie et al., 2016; Catizone et al., 2020). Our observations suggest the AP1 motif adjacent to the p53RE in the *GADD45A* intronic enhancer is required for maximal transcriptional output but is not required for p53-dependent induction. Conversely, this AP1 binding site is strictly required for tunicamycin-induced enhancer activity, suggesting that the regulatory potential of this enhancer is context-dependent. The *GADD45A* intron 3 enhancer is predicted to encode at least 10 distinct TF motifs (JASPAR 2022 (Castro-Mondragon et al., 2022)). To identify additional TF motifs regulating context-dependent enhancer activity, we measured basal and stimulus-dependent enhancer activity using STARRSeq (self-transcribing active regulatory region sequencing) (Muerdter et al., 2018). We performed nucleotide-resolution saturating mutagenesis with all possible substitutions or single nucleotide deletion at every position within the 250 bp enhancer (Fig. 8A). First, this library was transiently transfected into the HCT116 wild-type and p53 null cell lines to assess p53 dependence. Cells were also treated with DMSO, nutlin-3A, or tunicamycin to measure p53 or ISR-mediated activation of the *GADD45A* enhancer reporter library (Fig. 8B, S5). Overall, most single-nucleotide substitutions and deletions had little or no effect on basal or drug-induced activity (Fig. 8B, S5).

**Figure 8.**
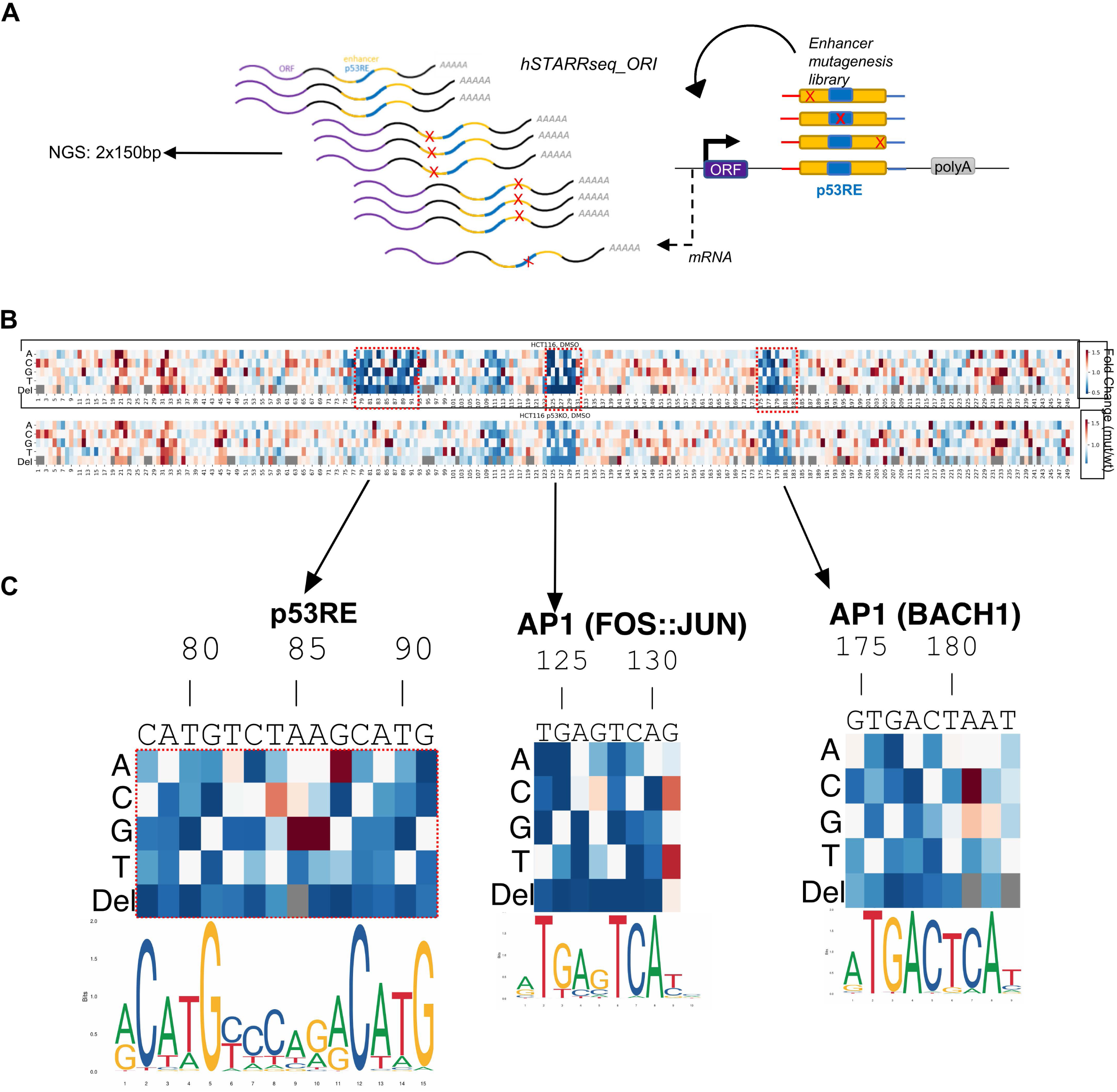
Nucleotide resolution of *GADD45A* enhancer sequence critical for promoter activation function. A) Schematic illustrating *GADD45A* 250 nt enhancer mutagenesis screen using the STARRSeq system (hSTARRSeq_ORI). Substitutions and deletions of every nucleotide position are depicted as ‘X’; p53RE in blue; open reading frame as ‘ORF’; polyadenylation site as ‘polyA’. B) Heatmap representing expression mediated by each enhancer variant relative to the wild-type enhancer from the same cell line and treatment condition. Relative position in the enhancer (1-250 nt) is indicated on the *x*-axis. Each deletion (‘Del’) or base substitution (‘A’, ‘C’, ‘G’, ‘T’) is indicated as a row label. ‘Grey’ color in row ‘Del’ indicates a redundant position when >1 consecutive base is identical. Relevant motifs discussed in the text are highlighted (red dashed line) including: *GADD45A* native motif sequence, relative position, name (‘p53RE’, ‘AP1’) and PWM logos (JASPAR 2022 (Castro-Mondragon et al., 2022)) are in panel C. Cell lines and treatment conditions are indicated above each heatmap.

Consistent with our *GADD45A-*full gene reporter assays (Fig. 7D), nucleotide substitutions and deletions at consensus motif positions with high predicted importance for p53 binding severely diminished enhancer activity in wild-type HCT116 cell line (Fig. 8C), with near perfect concordance to the canonical p53RE motif. Changes in the critical half-site positions of p53RE (nucleotide positions 78-81 and 88-91) consistently disrupted enhancer-driven transcription. Most nucleotide substitutions in the 6 bp spacer between the half-sites had minimal or no effect on enhancer activity (nucleotide positions 83-87). For example, T>A or T>C substitutions at the first position of the spacer are well-tolerated, but in agreement with known p53 binding preferences, T>G substitution reduced enhancer activity (Fig. 8C). T>C (position 3 of the spacer) and G>A (position 6) substitutions increased enhancer activity, again mirroring the shift closer to consensus p53 binding preferences. Conversely, all single nucleotide deletions disrupted enhancer activity, illustrating the well-studied importance of spacing between half-sites for p53 binding and activity (el-Deiry et al., 1992) (Fig. 8C). Comparison of nucleotide substitutions between WT and p53 null conditions suggests most of the substitutions with altered activity require an intact p53 response, except for A85G and A86G substitutions. Although these nucleotide changes are predicted to be more like the consensus sequence than wild type RE and display increased enhancer activity, they also show increased activity in p53 null cells, suggesting these substitutions may result in a *de novo* activating TF motif or disruption of a repressive TF element. Overall, loss-of-function substitutions in the p53RE did not display reduced enhancer activity in p53 null cells, as the genetic loss of p53 is expected to be epistatic with mutation of the p53RE (Fig. 8B). Nucleotide-resolution mutagenesis of the well-defined p53RE suggests that our STARRseq assay is suitable for high-throughput mutagenesis and identification of additional factors important for stress-dependent *GADD45A* enhancer.

We identified the AP1(FOS:JUN) site downstream of the p53RE, and not the predicted ATF4 upstream element, to be important for both basal– and tunicamycin-induced enhancer activity using traditional reporter gene assays (Fig. 7D). Disruption of the upstream ATF4 element in the saturating mutagenesis STARRseq assay had little to no effect on enhancer activity, whereas we observed a marked decrease in enhancer activity when the AP1 motif was disrupted in either WT or p53 null cell line (Fig. 8C, S5). The loss of activity in response to AP1 motif substitutions in p53 null lines further suggests combinatorial and additive roles of these two elements in driving basal enhancer activity. Specifiic substitutions to the AP1 (FOS:JUN) motif demonstrates a clear dependence for known AP1 family binding preferences on enhancer activity. C>G/T substitutions at position 131 have increased activity, presumably representing a shift closer to the consensus AP1 motif (Fig. 8C). The most sensitive positions to nucleotide substitutions were the two palindromic half-sites (‘TG’ and ‘CA’) and nucleotide deletions predicted to disrupt spacing between those half-sites (Fig. 8C).

Our STARRSeq assay for the *GADD45A* intron 3 enhancer revealed multiple nucleotide substitutions with increased or decreased activity. Individual nucleotide changes can disrupt TF motifs important for wild-type *GADD45A* enhancer activity but could also represent varied activity due to *de novo* creation of TF binding sites or experimental noise. Therefore, to identify other true positive TF motifs, we focused our analysis on contiguous 6+ nucleotide regions that display altered activity when (1) multiple substitutions and deletions at the same position have similar effects, (2) changes in several adjacent positions have similar effect, and (3) motif mutations have negative effect on the expression and thus are potentially bound by activators. Using such criteria, we identified 3 other regions that may positively regulate *GADD45A* enhancer activity. Each of these regions overlap a specific predicted TF motif, ETV6, AP1/BACH1 or GLIS3/POU6F (Fig. S5). AP1 and BACH1 are part of the broader bZIP family of DNA binding proteins and have highly similar motif preferences. Consistent with the p53RE and the AP1 (FOS:JUN) sites, nucleotide substitutions at positions predicted to be important for either AP1 or BACH1 binding display reduced enhancer activity (Fig. 8C). These effects are observed in both WT and p53 null cell lines, suggesting this motif contributes to basal activity in a p53-independent context. An A>C substitution at position 181 strongly increases enhancer activity, likely due to C adhering more closely to the predicted consensus binding motif for both AP1 and BACH1 dimers. Nucleotide deletions at any position reduced enhancer activity, whereas we observed a position-specific effect of substitutions mirroring consensus nucleotide preferences. Our saturating mutagenesis approach validates the critical role of the previously characterized p53 and AP1 binding sites in the positive regulation of the *GADD45A* enhancer, and revealed putative, novel TF motifs that may contribute to basal or induced enhancer activity.

### Validation of the STARRseq-defined effects of nucleotide substitutions within the GADD45A enhancer via a native-context reporter assay activity

To assess whether nucleotide substitutions in newly identified *GADD45A* enhancer motifs would have the same effect in a native gene context, we tested a series of ‘upregulating’ and ‘downregulating’ substitutions in our *GADD45A-nLuc* reporter system (Fig. 9). All *GADD45A*-*nLuc* enhancer mutants were transiently transfected into HCT116 wild-type and p53 null and treated with DMSO, nutlin-3A and tunicamycin. The AP1/BACH1(A181C) substitution moves this motif closer to the consensus sequence and is expected to improve binding and subsequent activation of *GADD45A*, whereas G177A would likely have the opposite effect. Indeed, using the native reporter system demonstrates that these substitutions behave as predicted from the MPRA-based approach (Fig. 8C, 9C). We next assessed how substitutions in the overlapping GLIS3 and POU6F2 motifs (Fig. S5) affect activity in the more native context. All nucleotide substitutions led to increased activity in the MPRA, suggesting these motifs and their bound factors may act to repress *GADD45A* transcription. In the native reporter system, substitutions in this overlapping GLIS3/POU6F motif increased enhancer activity (Fig. 9C), further suggesting that this sequence restrains activity of the *GADD45A* enhancer.

**Figure 9.**
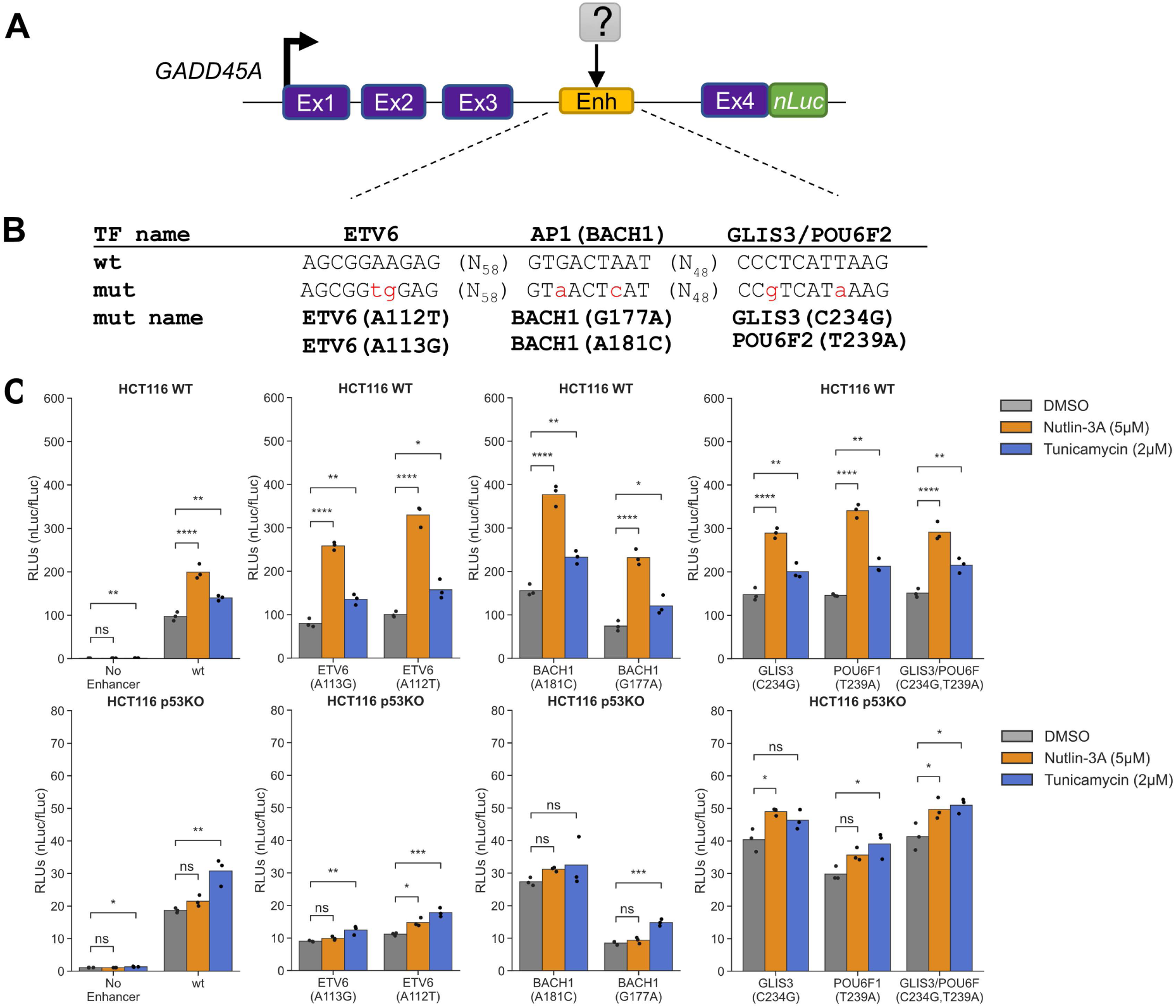
Other STARRSeq-identified motifs contribute only to basal *GADD45A* enhancer activity. A) Schematic representation of the *GADD45A-nLuc* reporter construct with relevant enhancer sequence motifs highlighted in panel B. C) Normalized luciferase expression values using *GADD45A-nLuc* reporter transfected into HCT116 WT cell line and 16h treatment with DMSO, nutlin-3A and tunicamycin as indicated in the legend. Reporter constructs included wild-type (wt) construct, 250 bp enhancer deletion (‘No Enhancer’) as a negative control and various predicted transcription factor motif mutations as indicated on the *x*-axis. Specific mutations in transcription factor motifs based on the STARRSeq screen are indicated in panel B. Statistical comparisons were generated using one-way ANOVA: * p<0.05, ** p<0.01, *** p< 0.001, **** p<0.0001.

Results from native reporter gene assays measuring the activity of nucleotide substitutions in the putative ETV6 motif located between the p53 and AP1 motifs were more nuanced (Fig. 9C). ETV6 is an ETS-family transcription factor and most frequently acts as a repressor (Lopez et al., 1999). The ETV6 A113G substitution, which led to consistently increased activity in the MPRA (Fig. S5), displayed increased nutlin-3A/p53-induced activity, but did not affect expression under basal or tunicamycin-treated conditions. The A112T substitution, predicted to reduce enhancer activity, surprisingly displayed higher levels of nutlin-3A-induced activity in HCT116 WT p53 using the native reporter system. Interestingly, both the ETV6 A112T and A113G substitutions led to diminished enhancer activity across all treatment conditions in HCT116 p53 null. These data suggest a potential context-dependence of the ETV6 motif in regulation of *GADD45A* enhancer activity, with motif disruptions leading to varied effects depending on the presence of p53.

## Discussion

In this study, we present a comparative analysis of gene regulatory strategies for two fundamental cell stress responses. We demonstrate the p53 gene regulatory network and the ATF4-driven Integrated Stress Response, although generally controlling distinct genes, converge on a set of common transcriptional targets related to metabolic control and apoptosis. Our study provides direct evidence that these targets require p53 during the DNA damage response, but not during the ISR. Conversely, ATF4 is required during the ISR and is dispensable under p53-activating conditions. The genetic dependence parallels the well-studied, stress-evoked stabilization of p53 and translation of ATF4 (Kastan et al., 1991; Vattem and Wek, 2004), which is further supported by observations that neither p53 nor ATF4 levels increase after activation of the other pathway (Fig. 1B, 2A, D). Importantly, these data are consistent with recent work demonstrating targeted ISR induction activates specific p53 gene targets in p53-deficient cells (Hernandez Borrero et al., 2021; Tian et al., 2021). Similarly, chemical inhibition of the phosphatase PPM1D leads to increased ATF4 activity which synergized with p53 to amplify expression of some p53 target genes and increased cell death (Andrysik et al., 2022). These data point towards therapeutic strategies broadly applicable across cancers regardless of *TP53* genetic status. Combined treatment with non-genotoxic activators of p53 (like MDM2 inhibitors) and chemical induction of the ISR pushes cells towards an apoptotic/cell death fate, which could overcome the reported limitations of MDM2 inhibition alone (Aziz et al., 2011). Small molecules restoring p53 function in p53-deficient tumors are being developed and their efficacy may be bolstered by combined ISR activation. These approaches are especially attractive given the non-genotoxic nature and the relatively large number of experimental and approved compounds that activate p53 or the ISR.

Both the p53-dependent and the ATF4-driven ISR gene networks are antiproliferative, either through induction of apoptosis or cell cycle control. p53 canonically mediates cell cycle control through *CDKN1A/*p21 and other targets, like *CCNG1* (Jensen, 2003). The p53 network appears to be “redundant”, where loss of a single antiproliferative strategy does not appreciably affect tumor suppressor function. These shared genes may represent an additional layer of redundancy to tumor suppression by p53 through metabolic control. At least four of the shared p53 and ATF4 targets (*DDIT4, GADD34, SESN2,* and *GDF15)* are antiproliferative via inhibition of mTOR signaling (Budanov and Karin, 2008; Gambardella et al., 2020; Lockhart et al., 2020; Aguilar-Recarte et al., 2021; Coronel et al., 2022). *ATF3* also coordinates cell cycle progression through serine, nucleotide, and glucose metabolic control (Ku and Cheng, 2020; Di Marcantonio et al., 2021). These genes, thus, may represent a “core” repurposed by various cell stress response pathways for anti-proliferative effects through the central energy regulator mTOR. Thus, investigations into stress-dependent regulation of these targets by other transcription factors controlling hypoxia (HIF1α), heat shock (HSF1), inflammation (IRF/STAT), xenobiotics (AHR), and infection (NF-kB), may be warranted.

Our genetic dissection of *ATF3* and *GADD45A* provides mechanistic detail into independent regulation by both p53 and ATF4. *ATF3* induction after stress is mediated by two spatially distinct and mutually independent regulatory elements, each bound by either p53 or ATF4. While our data suggest that basal *ATF3* expression is reduced when either element is perturbed, stress-mediated induction appears wholly dependent on the specific stress-activated transcription factor and its unique bound element. In contrast, stress dependent *GADD45A* induction is controlled by a single enhancer element bound by both p53 and ATF4. STARRSeq-based saturating mutagenesis of this *GADD45A* enhancer provided nucleotide-level resolution of DNA elements that control enhancer activity. Consistent with our genetic, biochemical, and reporter gene analyses, this assay demonstrated nucleotide substitutions at positions predicted to be critical for binding by either p53 or ATF4 most significantly impacted enhancer activity and *GADD45A* expression. This is true for predicted loss-of-function nucleotide substitutions, but also substitutions that are predicted to improve transcription factor binding (Fig. 8C). These data suggest that MPRA-style assays like STARRseq are suitable for examining the impact of sequence differences in p53-bound elements resulting from natural or disease-associated human variation (Fig. S5). This impact has not been comprehensively explored, but certain gain-of-function variants with pro-tumorigenic activity have been reported (Menendez et al., 2007; Zeron-Medina et al., 2013). We also identified and validated three additional transcription factor binding motifs directly impacting enhancer activity. Sequence-based motif prediction methods identify numerous putative transcription factor motifs within the *GADD45A* enhancer that did not alter enhancer activity in this context. Enhancers have well-defined cell lineage-dependent activity based on the specific combination of transcription factors that may be present (Zeitlinger, 2020; Kim et al., 2021), thus, we cannot rule out that these predicted motifs may be functional in other settings. Thus, a combination of unbiased saturating mutagenesis screening across diverse cell types and sequence-based motif analyses may help to refine and improve the ability to predict functional elements within regulatory elements.

Our results examining p53 and ATF4-mediated induction of *GADD45A* provide new insight into the complex interplay between multiple transcription factors at stress-dependent enhancers. The p53RE is critical for nutlin-3A-induced expression of *GADD45A,* whereas other motifs were dispensable, including the AP1 site critical for ATF4-mediated expression. These motifs are also unnecessary for ATF4-dependent induction. Our observations mirror prior studies demonstrating p53-induced enhancer activity solely depends on p53 binding and that p53 motifs are the strongest predictor of high enhancer activity (Verfaillie et al., 2016; Younger and Rinn, 2017). While global MPRA analyses of ATF4-dependent regulatory elements have not been reported, our results suggest that like p53, ATF4 may not require other transcription factors to induce stress-dependent enhancer activity. This model is attractive as the regulatory activity of stress-dependent transcription factors would remain robust across contexts. Although they appear unnecessary for stress-mediated induction, transcription factor motifs (ETV6, AP1:BACH1, GLIS3/POU6F) identified via saturating mutagenesis modulate *GADD45A* enhancer activity in unstressed conditions. These results also provide a context where additional transcription factors regulate p53– or ATF4-bound enhancers and suggest that differential enhancer “grammar” may be needed for basal versus stress-dependent transcriptional regulation.

p53-dependent enhancers may rely on other transcription factors in certain contexts, like with varied cell lineage-dependent chromatin structure (Karsli Uzunbas et al., 2019). One caveat to MPRA studies is they can lack chromatin context that might mask the requirement for other transcription factors that alter chromatin accessibility. Additionally, enhancers display cell lineage-dependent activity due to differential expression of these transcription factors and cofactors. These caveats for MPRA studies are also relevant for our identification of shared gene targets between the p53 and ISR gene networks. Our global analyses was limited to one colon carcinoma cell line, but recent studies have identified previously underappreciated cell lineage-dependent p53 transcriptional networks (Andrysik et al., 2017; Nguyen et al., 2018; Fischer, 2019; Karsli Uzunbas et al., 2019). A full analysis of cell lineage-specificity has not been directly explored in the ATF4-dependent ISR. Thus, the shared gene network between the p53 and ISR gene networks is likely more expansive than is reported here. Our results identify autonomous activity of p53 and ATF4 during stress-dependent enhancer activation and gene regulation, and certainly additional exploration of context-dependent enhancer activity and shared gene targets is required,

## Data Availability

The data that support the findings of this study are openly available in Gene Expression Omnibus (GEO) at https://www.ncbi.nlm.nih.gov/geo/, reference numbers GSE211244 (RNAseq), GSE211264

(CUT&RUN), GSE226617 (STARRSeq mutagenesis screen).

## Acknowledgments

The authors would like to thank the University at Albany Center for Functional Genomics for sequencing and analytical support. SAD was supported by a University at Albany RNA Institute Fellowship. This work was supported by NIH NIGMS R35 GM138120.

---

**Figure S1.**
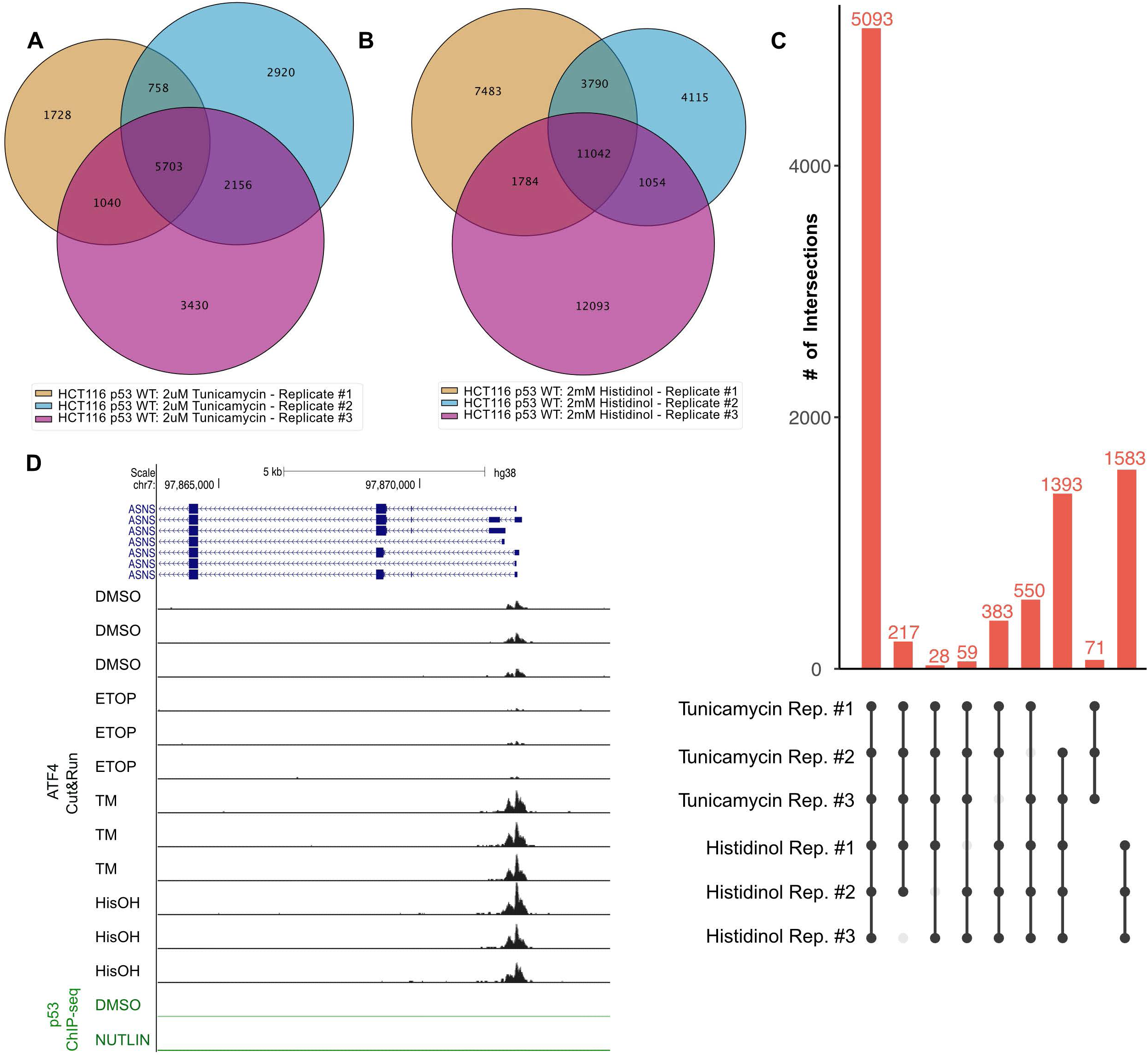
A) Intersection of ATF4 CUT&RUN peaks for the three biological replicates of HCT116 p53 WT cells treated with 2 μM tunicamycin for 6h. B) Intersection of ATF4 CUT&RUN peaks for the three biological replicates of HCT116 p53 WT cells treated with 2 mM histidinol for 6 hrs. C) A set of high-confidence ISR-activated ATF4 binding events created by considering only peaks called out from 5 out of the 6 experiments with ISR-activating treatments: 2 μM tuncamycin and 2 mM histidinol. D) Genome browser view of the *ASNS* gene locus displaying ATF4 CUT&RUN data (black) and p53 ChIP-Seq data (green) in HCT116 p53 WT cells following 6 h treatment with various stress stimuli, including: DMSO (vehicle control), 5 μM nutlin-3A (NUTLIN), 100 μM etoposide (ETOP), 2 μM tunicamycin (TM), and 2mM histidinol (HisOH).

**Figure S2.**
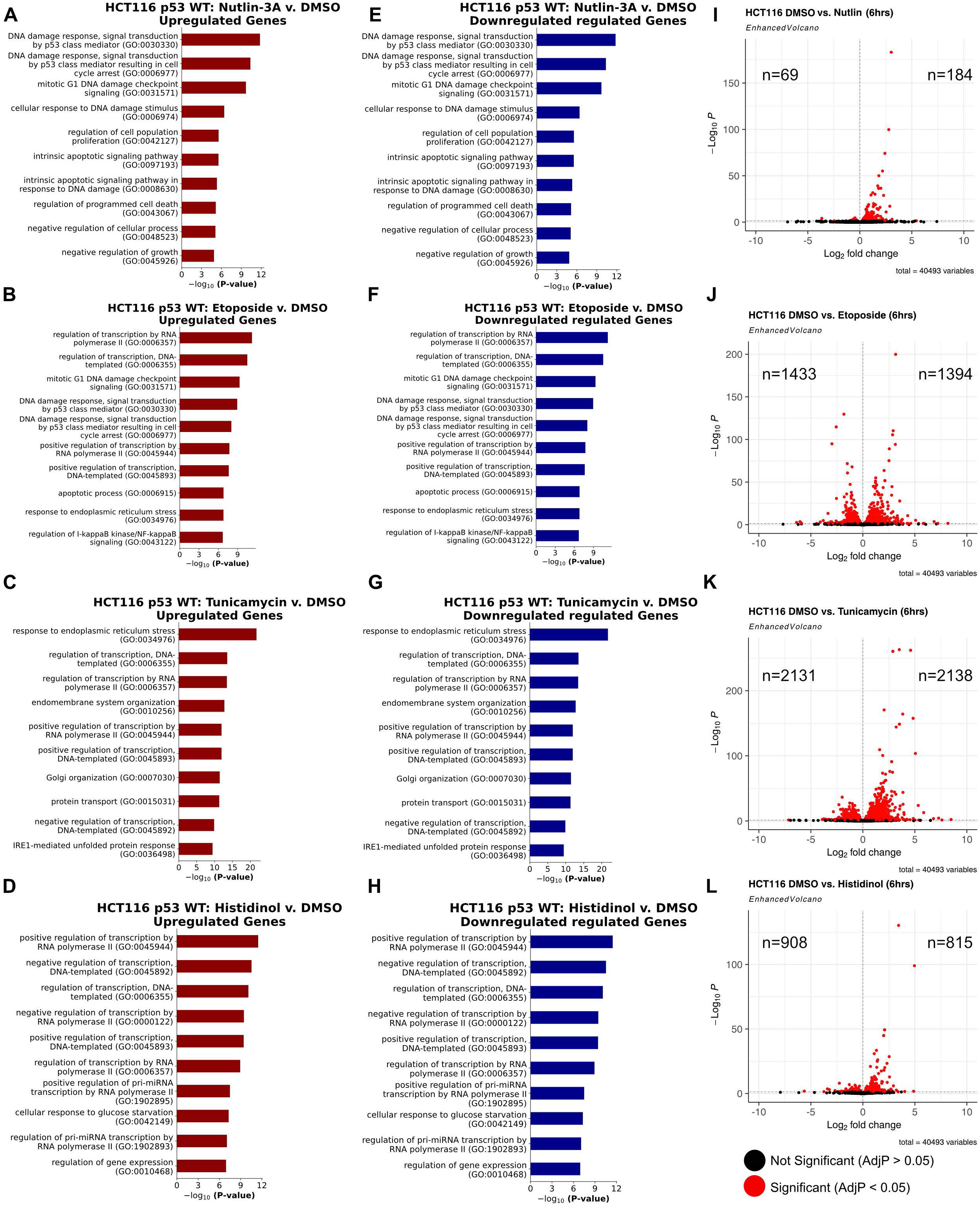
RNASeq analysis of differential gene expression in HCT116 WT and p53 null cell lines in response 5 μM nutlin-3A (A, E), 100 μM etoposide (B, F), 2 μM tunicamycin (C, G), or 2mM histidinol (D, H), compared to vehicle control (DMSO). Gene ontology analysis of the genes upregulated (A-D) and downregulated (E-H) in response to treatments. (I-L) Enhanced volcano plots displaying differential gene expression in HCT116 WT cells treated stimuli as described above. Number of genes (n) upregulated or downregulated in each condition is indicated on the left side or the right side of each volcano plot, respectively.

**Figure S3.**
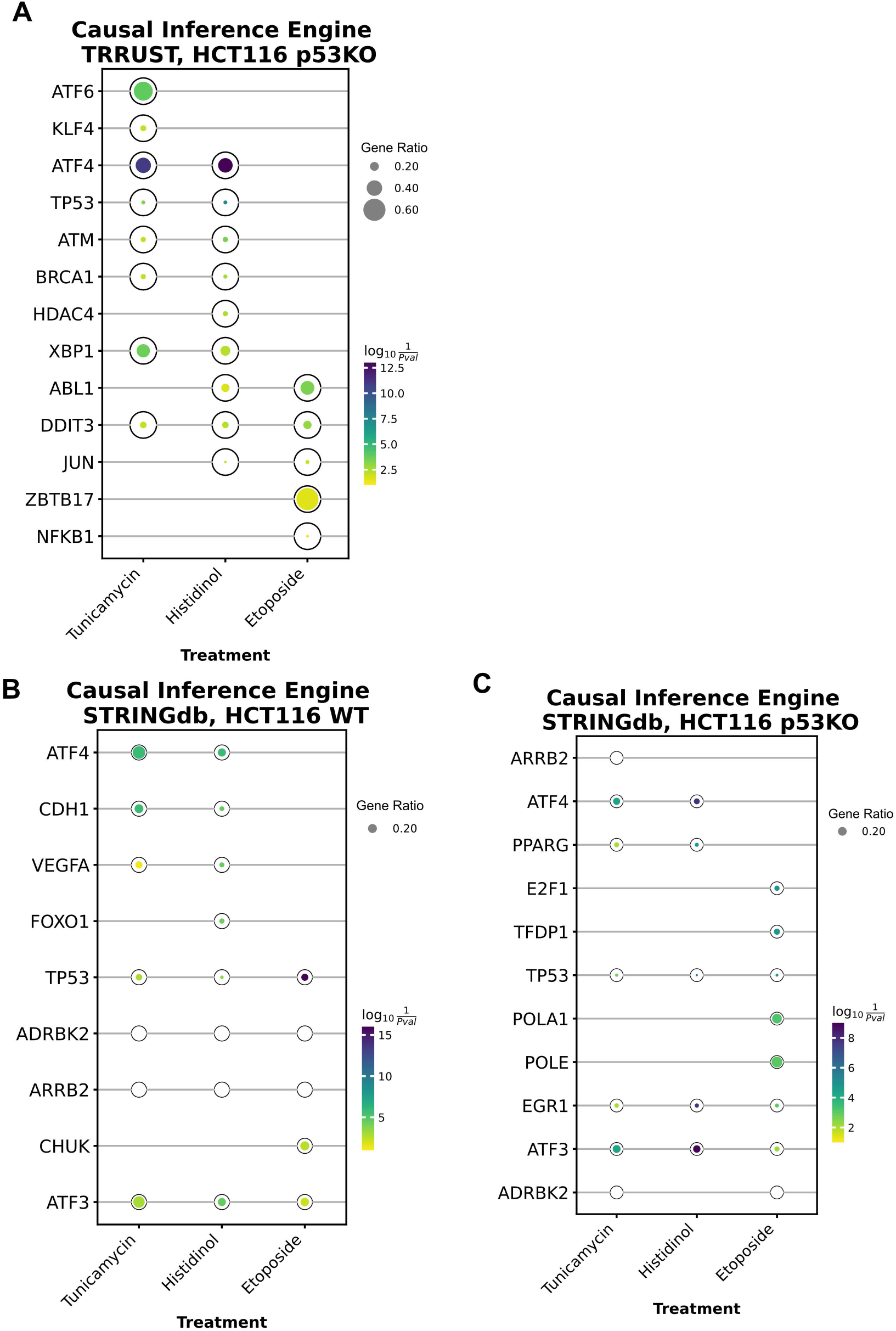
Causal Inference Engine enrichment of putative upstream regulators of stress-induced gene expression in WT and p53 null HCT116 cells. A) Results of the top 5 most enriched upstream regulators from the TRRUST database of tunicamycin, histidinol, or etoposide-induced genes in HCT116 p53 null cells (p-value derived from the Fisher’s Exact test for enrichment). Results of the top 5 most enriched upstream regulators from the STRING database for genes induced in (B) HCT116 WT or (C) HCT116 p53 null cells from Causal Inference Engine (p-value derived from the Quaternary Scoring statistic (QS) from (Fakhry et al., 2016)). Plots were generated with GSEApy (Fang et al., 2023).

**Figure S4.**
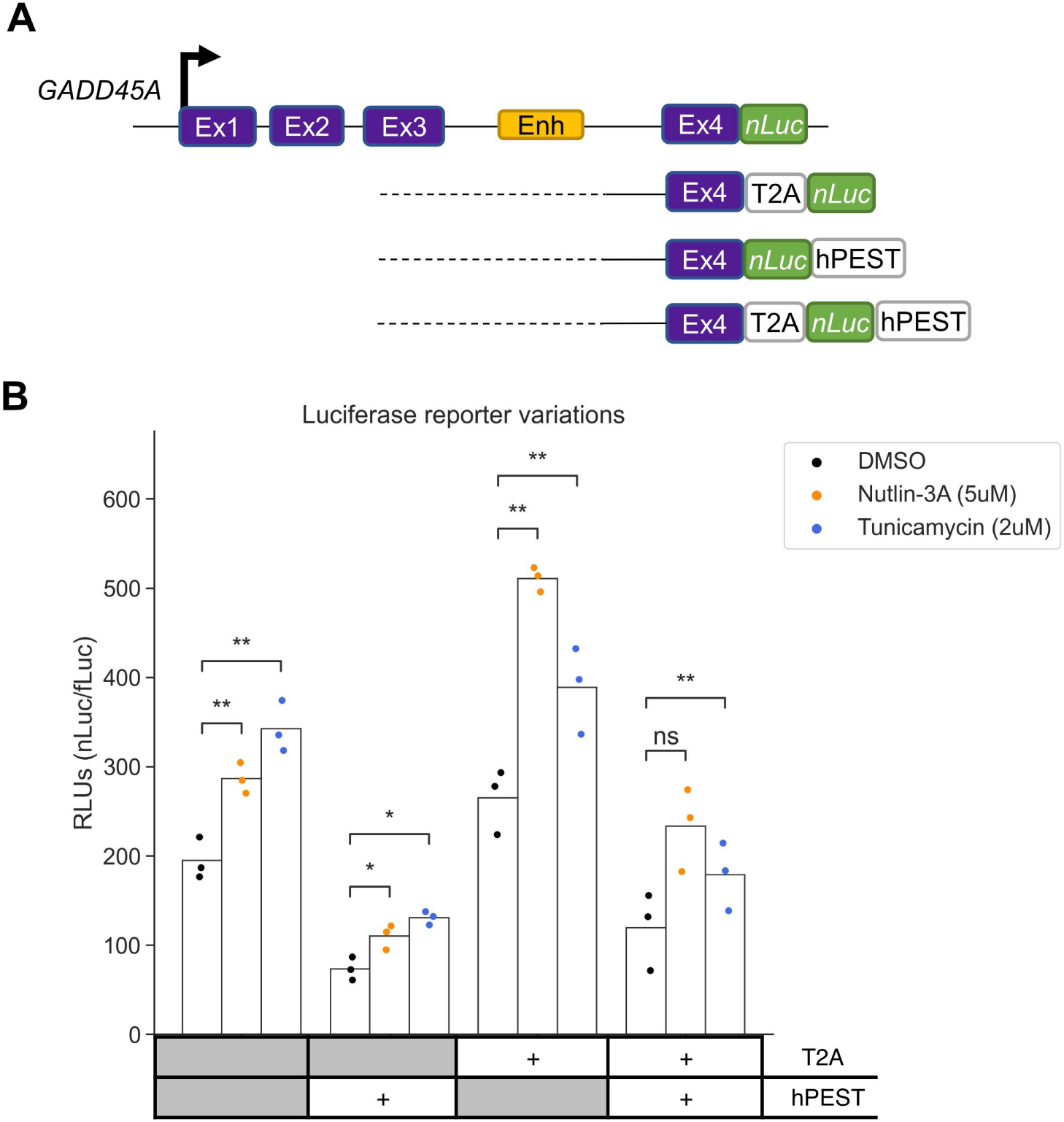
Comparison of *GADD45A-nLuc* reporter expression in the presence or absence of T2A skipping peptide and/or hPEST degradation tag. A) Schematic representation of various *GADD45A–nLuc* reporter constructs with T2A or hPEST included in the 3’ end. B) Normalized luciferase expression values from *GADD45A-nLuc* reporters transfected into HCT116 WT cell line and 16h treatment with DMSO, nutlin-3A and tunicamycin as indicated in the legend. Presence (‘+’) or absence (grey) of T2A or hPEST tags are indicated in the table below.

**Figure S5.**
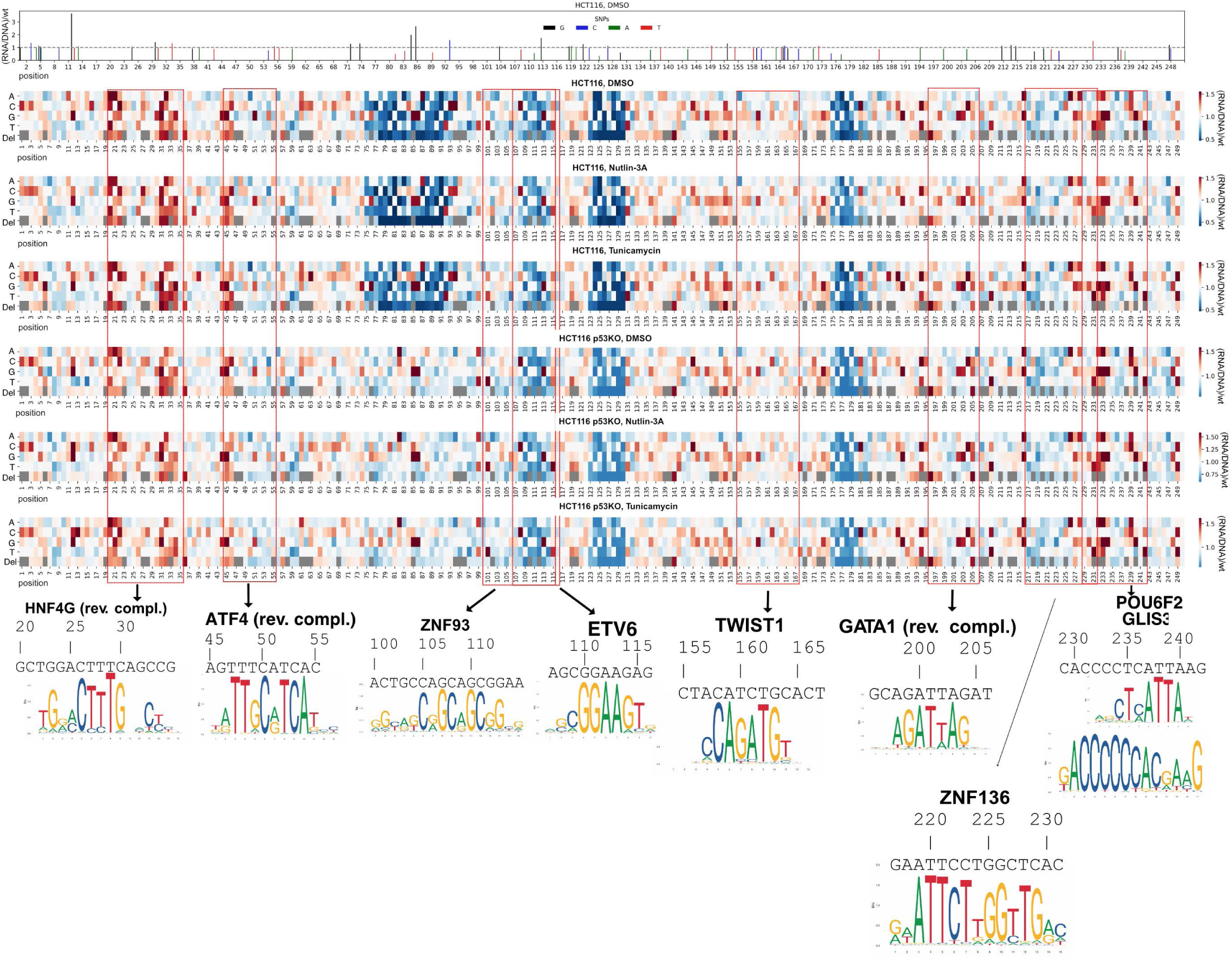
Nucleotide resolution of *GADD45A* enhancer sequence critical for promoter activation function (extended Fig. 8B). Heatmap representing expression mediated by each enhancer variant relative to the wild-type enhancer from the same cell line and treatment condition. Relative position in the enhancer (1-250 nt) is indicated on the *x*-axis. Each deletion (‘Del’) or base substitution (‘A’, ‘C’, ‘G’, ‘T’) is indicated as a row label. ‘Grey’ color in row ‘Del’ indicates a redundant position when >1 consecutive base is identical. Relevant motifs discussed in the text are highlighted (red dashed line) including: *GADD45A* native motif sequence, relative position, name (’HNF4G’, ‘ATF4’, ‘ZNF93’, ‘ETV6’, ‘TWIST1’, ‘GATA1’, ‘ZNF136’, ‘GLIS3/POU6F2’) and PWM logos (JASPAR 2022 (Castro-Mondragon et al., 2022)). Cell lines and treatment conditions are indicated above each heatmap. (Top Row) Barplot highlighting SNPs found in *GADD45A* enhancer region from db155SNP database with expression values from HCT116 WT, DMSO experiment.

Table S1. List of plasmids and oligonucleotides used in this study.

Table S2. Full list of DEseq2-derived differential gene expression values for HCT116 WT and p53 null cells treated with vehicle control (DMSO) relative to nutlin-3A, etoposide, tunicamycin, or histidinol treatments.

Table S3. Full Gene Ontology Enrichment results from Figure 5 and Figure S2.

Table S4. Complete Causal Inference Engine analysis of putative upstream regulators of genes induced by etoposide, tunicamycin, or histidinol in WT or p53 null HCT116 cells.

Table S5. Full *chipenrich* results using KEGG, HALLMARK and REACTOME databases.

